# Differential Adaptation of Osteocyte Lacunar Morphology at Bone (Re)modeling Sites under Mechanical Loading and Pharmacological Therapies in a Mouse Model of Osteoporosis

**DOI:** 10.64898/2026.10.05.756707

**Authors:** Sara Lindenmann, Zihui Li, Claudia Weigt, Floor M. Lambers, Friederike A. Schulte, Gisela A. Kuhn, Neashan Mathavan, Ralph Müller

## Abstract

Postmenopausal osteoporosis is a highly prevalent disease characterized by low bone mass and structural deterioration of bone tissue leading to increased bone fragility and fracture risk. Osteocytes orchestrate local bone (re)modeling and therefore, characterizing their 3D morphological and spatial distribution across distinct (re)modeling microenvironments is essential for understanding the cellular mechanisms underlying divergent tissue-level responses. To address this, we used a 3D-3D multiscale registration framework mapping endpoint osteocyte lacunae onto dynamic, time-lapsed *in vivo (*re)modeling sites. Ovariectomized 15-week-old mice were subjected to 4 weeks of anabolic parathyroid hormone (PTH) or anti-catabolic bisphosphonate (BIS) treatment as monotreatment or combined with cyclic mechanical loading (ML). Biweekly *in vivo* micro-computed tomography (micro-CT) to identify (re)modeling sites and endpoint ultra-high resolution (ultra-HR) micro-CT to characterize osteocyte lacunae was performed. Osteocyte lacunae were characterized by lacunar density, volume and sphericity. Our findings showed that physical and pharmacological therapies differentially regulate the osteocyte lacunar network through distinct perilacunar adaptations across functional microenvironments. While BIS stabilizes local lacunar architecture, PTH drives asymmetrical lacunar expansion (increased volume with reduced sphericity) through perilacunar remodeling that supersedes any added ML effects. Crucially, estrogen loss disrupts the natural spatial maturation gradient by inducing rounding of lacunae in newly formed bone. Therapies counteract this disruption, restoring the healthy gradients in lacunar morphometry between spatially distinct microenvironments. Additionally, despite restoring tissue-level bone volume, physical and pharmacological osteoporosis therapies do not fully recover persistent lacunar density deficits. The overall lacunar morphological spatial signatures confirm active osteocytic tuning to endocrine, physical and matrix cues.

## 1. Introduction

In the 19^th^ century, Julius Wolff^1^ formulated the theory that bone is a living tissue adapting to functional and mechanical requirements. During the same era, Roux^2^ proposed that this adaptive response to external loads is regulated by cellular activity. By the early 21^st^ century, osteocytes were identified as the primary mechanosensors within bone, mediating its adaptive response to external mechanical cues.^3,4^ Osteocytes lie within lacunae embedded in the bone matrix (extracellular matrix or ECM) and are connected through dendritic processes in the surrounding canaliculi. Osteocytes actively modify the matrix immediately surrounding their lacunae, termed the perilacunar matrix (PLM). Together, osteocytes and their dendritic processes form an elaborate lacuna-canalicular network (LCN) through which they sense mechanical loading via direct interactions to the surrounding tissue and fluid-flow-induced shear stresses.^5–8^ Beyond mechanosensation, osteocytes can actively maintain and adapt their immediate pericellular matrix through osteocytic perilacunar remodeling (PLR).^9^

The response and efficiency of this mechanosensory network to mechanical stimuli depend on osteocyte spatial density, dendritic connections and 3D lacunar morphometry.^10^ While osteocyte density is closely linked to bone (re)modeling rate,^3^ cellular shape and local 3D lacunar geometry directly dictate mechanosensitive strain amplification,^10,11^ making local lacunar structure a crucial determinant of cellular responsiveness across distinct bone microenvironments.

Bone disorders such as osteoporosis, osteopetrosis and osteogenesis imperfecta exhibit distinct deviations from healthy osteocyte lacunar morphology and distribution.^12–16^ It has been shown that changes in lacunar morphology coincide with changed mechanosensitivity where rounder morphology indicates higher mechanosensitivity.^17^ Consequently, increasing attention has been directed toward characterizing the osteocyte network across healthy and diseased states. Postmenopausal osteoporosis is a highly prevalent disease characterized by low bone mass and structural deterioration of bone tissue leading to heightened fracture risk^18^. Estrogen loss not only enhances osteoclast-mediated bone resorption, but also increases osteocyte apoptosis,^19,20^ resulting in reduced osteocyte density in both humans^21,22^ and rodents.^23^ Current therapeutic strategies rely on anti-resorptive and anabolic agents to specifically regulate the population and activity of osteoblasts and osteoclasts to counteract bone loss. Anti-resorptive bisphosphonates (BIS) inhibit bone resorption by binding to hydroxyapatite crystals in bone, thereby disrupting osteoclast attachment and activity.^24^ Nitrogen-containing BIS, such as zolendronic acid, further dysregulate core osteoclast cellular activities to induce osteoclast apoptosis ^25^. Anabolic parathyroid hormone (PTH) analogs stimulate bone formation by activation of osteoblasts. Interestingly, both BIS and PTH show secondary anti-apoptotic effects on osteocytes through distinct molecular pathways.^26,27^ In parallel, mechanical loading activates osteocytic mechanotransduction to drive anabolic bone adaptation.^27–30^ Furthermore, the mechanical environment shapes osteocyte morphology through cytoskeletal attachments to the pericellular matrix.

While the individual pathways of these pharmacological and mechanical stimuli are well-documented, how they interact at the cell scale to regulate bone (re)modeling through osteocytes remains poorly understood. Recently, Schulte et al. demonstrated that combining mechanical loading with PTH results in an additive effect in predicted bone strength, whereas combining mechanical loading with BIS offers no therapeutic advantage.^31^ However, while these macroscale functional outcomes are promising, the underlying cellular adaptations remain unknown. Because osteocytes orchestrate local bone (re)modeling, characterizing their 3D morphology and spatial distribution across distinct (re)modeling microenvironments under combined pharmacological and mechanical stimulation is essential to deciphering these divergent tissue-level responses.

Evaluating osteocyte lacunae within their distinct (re)modeling microenvironments poses a major technical challenge. To unravel these localized cellular adaptations, multiscale correlative mapping of time-lapsed *in vivo* micro-computed tomography (micro-CT) onto ultra-high-resolution (ultra-HR) *ex vivo* micro-CT images offers a powerful solution. Time-lapsed *in vivo* micro-CT allow tracking of specific sites of bone formation, resorption and quiescence over time while endpoint ultra-HR micro-CT allows accurate characterization of individual osteocyte lacunae. To address this challenge, this study established a 3D-3D multiscale registration framework to map endpoint osteocyte lacunae onto dynamic, time-lapsed *in vivo (*re)modeling sites in a mouse model of postmenopausal osteoporosis (Fig. 1). Using this pipeline, we investigated how pharmacological therapies (PTH, BIS) and mechanical loading uniquely alter osteocyte spatial distribution and 3D lacunar morphology both globally and within localized (re)modeling microenvironments. Based on previous tissue level findings,^31^ we hypothesized that anabolic PTH dynamically enhances osteocyte pericellular (re)modeling to facilitate a coordinated, synergistic tissue adaptation when combined with mechanical loading. Conversely, we hypothesized that the anti-catabolic mechanism of BIS extends to the cellular scale, yielding a stabilized, unreactive osteocyte network that derives no additional morphological benefit from mechanical stimulation. Finally, we hypothesize that osteocyte lacunar distribution and 3D morphology varies systematically between distinct (re)modeling microenvironments, with therapeutic effects reflecting depth-dependent mechanisms: systemic PTH drives widespread PLR extending into the quiescent bone whereas BIS selectively modulates surface-adjacent and thus mostly newly formed bone. Ultimately, this work establishes a validated multiscale methodology linking local remodeling dynamics to osteocyte morphometry, providing unprecedented spatial insight into osteocyte morphology and spatial organization in response to osteoporosis treatments.

**Figure 1:**
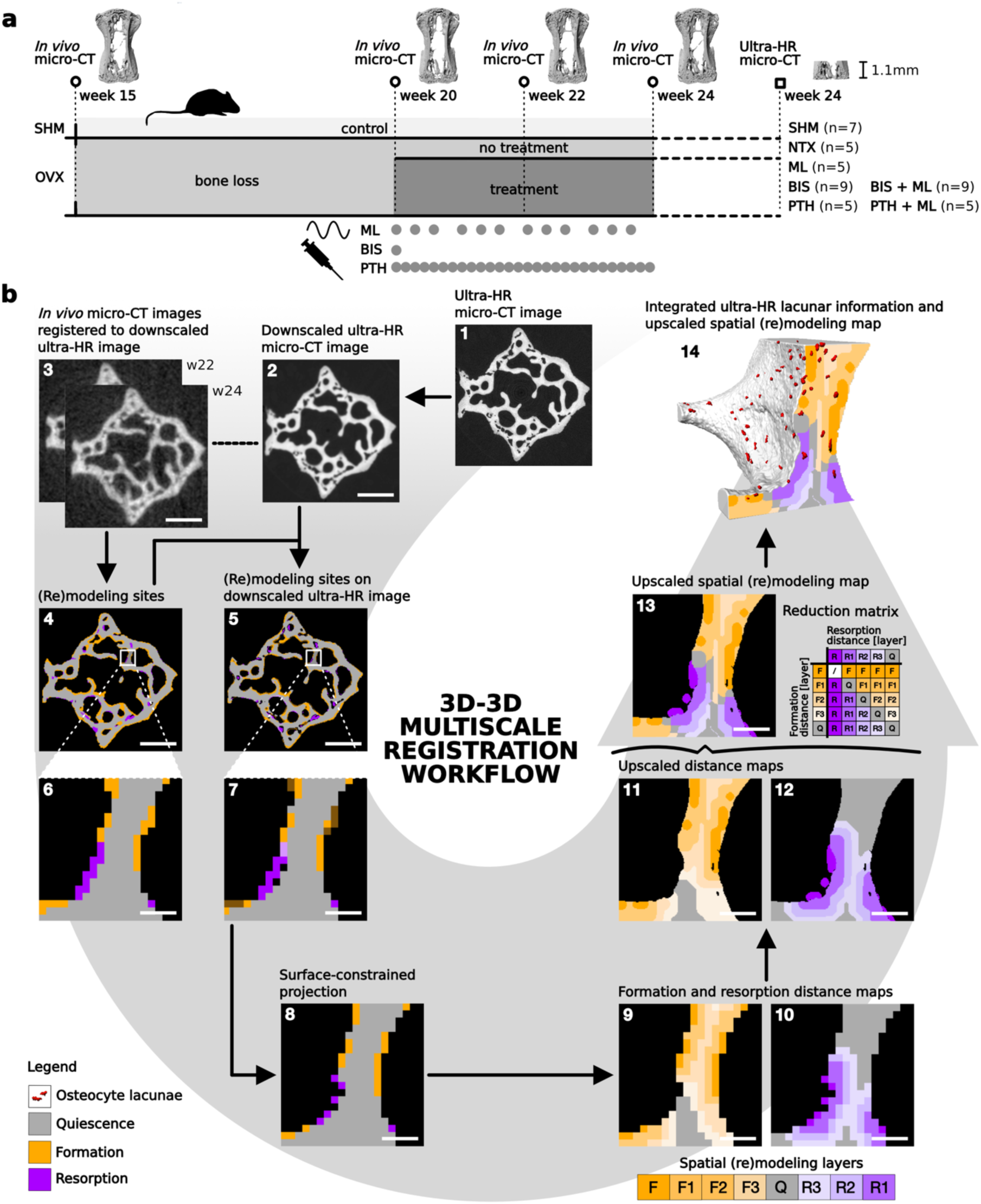
3D-3D multiscale registration of micro-computed tomography (micro-CT) images allows investigation of spatial differences in osteocyte lacunar morphology. (a) Experimental setup. 15-week-old mice were ovariectomized (OVX) or underwent sham surgery (SHM), and treatment was administered starting at week 20 (parathyroid hormone (PTH), bisphosphonates (BIS) and mechanical loading (ML)). Time-lapsed in vivo micro-CT images were obtained at week 15, 20, 22 and 24, and ultra-high-resolution (ultra-HR) ex vivo micro-CT images were obtained 10 hours after sample extraction. (B) Visual overview of the 3D-3D multiscale registration workflow starting with the ultra-HR micro-CT image (b1) before and (b2) after downscaling. (b3) The week 22 (w22) and week 24 (w24) in vivo micro-CT images were registered to each other and subsequently registered to the downscaled ultra-HR micro-CT image. (b4-7) Bone formation and resorption volumes were identified on the segmented overlays of the in vivo micro-CT images and applied to the ex vivo micro-CT bone volume. (b8) A surface constrained projection was applied to the (re)modeling volumes. (b9-12) Formation and resorption distance maps were created and upscaled in the following spatial (re)modeling layers: formation (F), formation adjacent layers 1-3 (F1, F2, F3) quiescent bone (Q), resorption adjacent layers 1-3 (R1, R2, R3). (b13) The ultra-HR distance maps were combined to one spatial (re)modeling map using the indicated reduction matrix. (b14) Integration of the spatial (re)modeling and the lacunar map for spatial analysis of osteocyte lacunar morphology. Scale bars: 50 µm.

## 2. Results

### Pharmacological and mechanical therapies reverse ovariectomy-induced bone loss

Trabecular bone loss in ovariectomized mice (n = 42) was confirmed by *in vivo* micro-CT of the whole 6^th^ caudal vertebra from week 15 to 20 (−23.9 ± 10.1% change in bone volume fraction (BV/TV) compared to baseline, data presented as mean ± standard deviation). On the other hand, the sham (SHM)-group showed a growth-related increase in trabecular bone over the initial five weeks of the experiment (16.8 ± 7.3% change in BV/TV). From week 20 to 24, all animals received either no treatment or pharmacological and/or mechanical treatment through an established tail vertebra loading model^32^. In that time interval, the trabecular BV/TV continued to decrease in the no treatment (NTX)-group while significantly increasing in all mono- and combined treatments, with the highest increase in PTH + ML (p < 0.01, Fig. 2b, Supplementary Table 3). BIS monotreatment showed the lowest increase in BV/TV and a non-significant difference to the SHM- and NTX-group (Fig. 2b, Supplementary Table 3). BV/TV of the SHM-group stayed constant over time (p > 0.05). The change in bone surface per total volume (ΔBS/TV) did not differ significantly from baseline in the SHM- and mechanically loaded (ML)-group but was significantly decreased compared to baseline in the NTX-group (p < 0.01, Fig. 2c). Only pharmacological mono- or combined treatment showed significantly increased ΔBS/TV at week 24 (Fig. 2c, Supplementary Table 3). Trabecular thickness (Tb.Th) remained unchanged in the SHM-, NTX- and BIS-group. PTH and ML increased trabecular thickness significantly over four weeks (p < 0.01, Fig. 2d). Trabecular number (Tb.N) showed a significant decrease in the NTX and ML-group as well as a small but significant decrease in the PTH-group (p < 0.05, Fig. 2e). As described by Schulte et al.^33^, BIS and PTH both increased BV/TV mainly through preserving existing trabeculae and minimal thickening for BIS and moderate thickening for PTH treatment. Pharmacological treatment in combination with ML did not change the trabecular number but significantly increased trabecular thickness compared to the pharmacological mono-treatment (p < 0.001, Supplementary Table 3). A full summary of these microstructural morphometric parameters is provided in the Supplementary Table 3.

**Figure 2:**
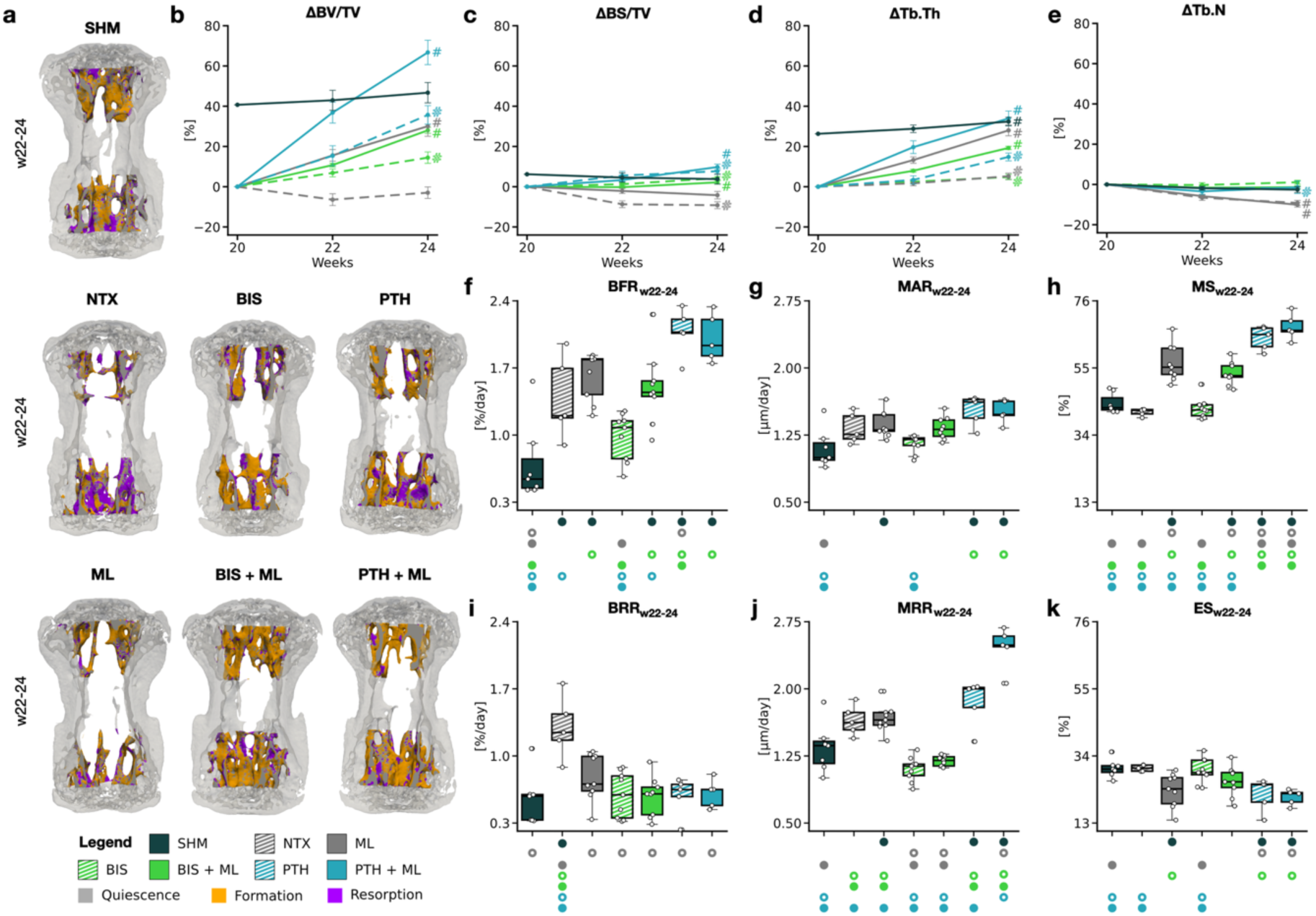
Static and dynamic morphometric parameters in trabecular bone from in vivo micro-computed tomography (micro-CT). (a) Representative micro-CT overlays at interval w22-24 with (re)modeling volumes. (b-e) Line plots showing the change in parameter in % compared to week 20. (b) Bone volume fraction (ΔBV/TV), (c) bone surface per total volume (ΔBS/TV), (d) trabecular thickness (ΔTb.Th) and (e) trabecular number (ΔTb.N). # indicates that the delta of the group from week 20 to 24 is different from zero as tested with t-test with null-hypothesis. (f-k) Boxplots representing the dynamic morphometric parameter between w22-24 for all groups. F) Bone formation rate per bone volume (BFR), (g) mineral apposition rate (MAR), (h) mineralizing surface per bone surface (MS), (i) bone resorption rate per bone volume (BRR), (j) mineral resorption rate (MRR) and (k) eroded surface per bone surface (ES). Significances were calculated using ANOVA followed by Tukey’s post-hoc where significant differences are depicted with a symbol in the color of the corresponding group. (o: to monotreatment, ●: to combined treatment, p < 0.05).

The time course of formation and resorption led to the observed changes in static morphometry described above (Fig. 2a). Bone formation rate per bone volume (BFR/BV) during week 22 to 24 (see also Supplementary Figure 4 for results from the first interval w20-22) was significantly increased in NTX compared to SHM (p < 0.05, Fig. 2f). Similarly, BFR/BV was significantly increased for ML, BIS + ML, PTH and PTH + ML (p < 0.001). PTH and PTH + ML additionally showed higher BFR/BV compared with the other treatments, consistent with the anabolic effect of PTH. Bone resorption rate per bone volume (BRR/BV) was significantly increased in NTX compared to SHM (p < 0.001, Fig. 2i) but not significantly different for any treatment. The mineral apposition rate (MAR) and resorption rate (MRR) did not differ significantly before and after ovariectomy (Fig. 2g and 2j). While MAR stayed relatively constant and didn’t show significant differences between NTX and any additional treatment, MRR significantly decreased for BIS and BIS + ML (p < 0.001) and significantly increased for PTH + ML (p < 0.001). The mineralizing surface (MS) and eroding surface (ES) showed no significant change with ovariectomy (Fig. 2h and 2k). Additional mechanical loading, however, resulted in significantly higher MS (p < 0.001) and lower ES (p = 0.057) compared to the NTX-group. BIS mono-treatment showed no significant change in MS and ES but additional ML resulted in a significantly higher MS than without (p < 0.001). Similarly, PTH treatment significantly increased MS (p < 0.001) and significantly decreased ES (p > 0.05) compared to NTX. A full summary of the dynamic morphometric parameters and statistical significances is provided in Supplementary Table 4.

### Physical and pharmacological interventions distinctly reshape global osteocyte lacunar morphology

Next, we investigated whether trabecular BV/TV in the 1.1 mm thick section of the ultra-HR micro-CT shows ovariectomy-induced bone loss observed on the time-lapsed *in vivo* micro-CT images. Ovariectomy resulted in a significantly lower endpoint BV/TV in the NTX-group compared to the SHM-group (Table 1, Fig. 3c). Pharmacological treatments and ML as mono-treatments or in combination showed a trend of increased BV/TV, with a significantly higher BV/TV in the PTH + ML-group than the NTX-, ML-, BIS-and BIS + ML-groups. The BIS-group furthermore shows significantly lower ΔBV/TV than the SHM-group. Importantly, endpoint sub-section results (Fig. 3c) agree with the longitudinal BV/TV observations observed for the whole trabecular bone from *in vivo* micro-CT images (Fig. 2b). The lacunar number (Lc.N) was significantly lower in all ovariectomized groups compared to the SHM-group (Table 1, Fig. 3d). The SHM-group also showed notably higher variability, both visually and quantitatively with a coefficient of variation (CV) of 25.3% while ovariectomized groups showed a CV of 10.6-23.3% (Table 1). This indicates that the number of lacunae in the samples of the SHM-group were more dispersed relative to the group mean while other groups appeared more consistent. Furthermore, lacunar density (Lc.N/BV), integrating both BV/TV and Lc.N, was significantly higher in the SHM-group compared to all other ovariectomized groups (Table 1, Fig. 3e). The ovariectomy-induced drop in lacunar density could not be recovered by either mono-or combined-treatment even though an increase in bone volume fraction could be observed.

**Table 1:**
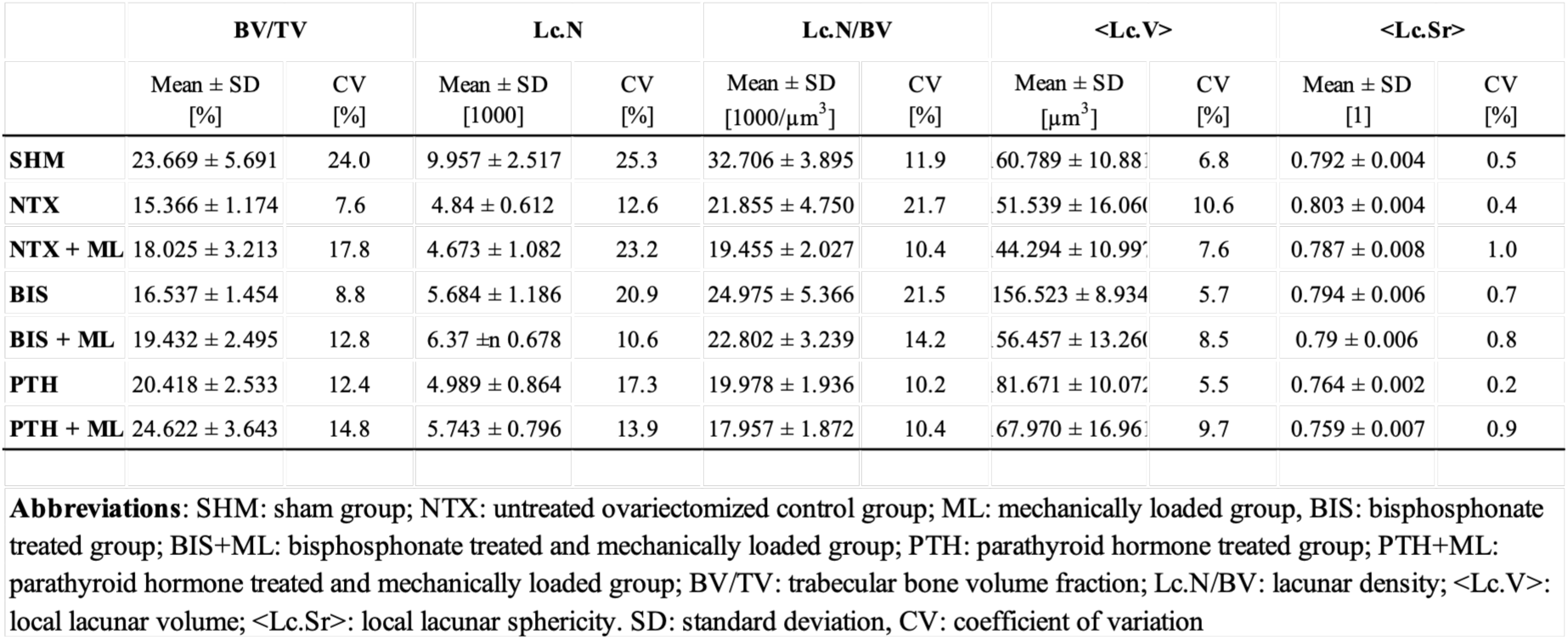
Mean, standard deviation (SD) and coefficient of variation (CV) for bone volume fraction (BV/TV) and lacunar distribution and morphology (Lc.N, Lc.N/BV, <Lc.V> and <Lc.Sr>) across groups calculated from the ultra-high-resolution (ultra-HR) micro-CT images.

**Figure 3:**
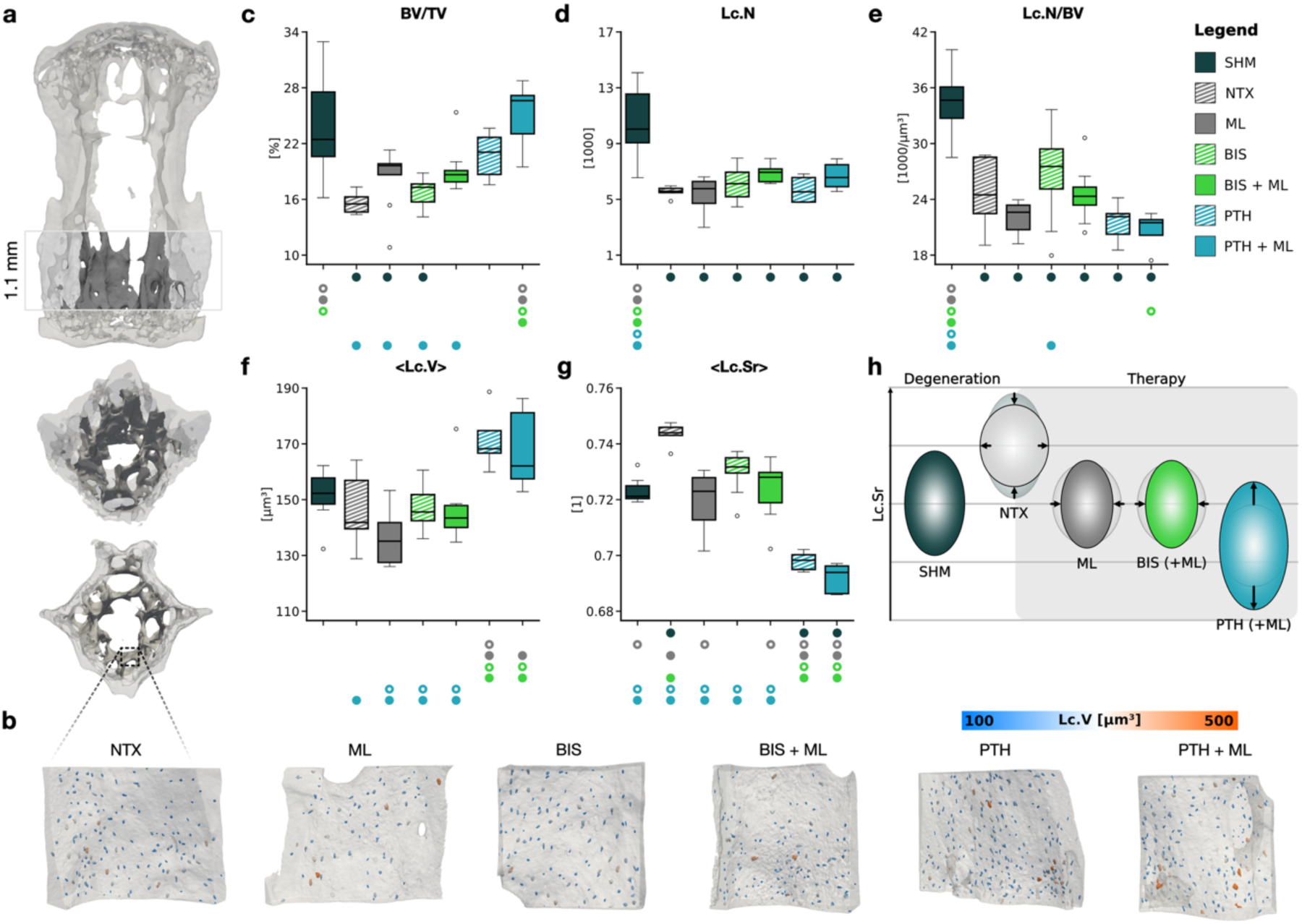
Global lacunar morphometry in the ultra-high-resolution (ultra-HR) micro-computed tomography (micro-CT) section of trabecular bone compared between all groups. (a) Representative in vivo micro-CT with extracted section scanned in ultra-high resolution. (b) 3D orthographic projections of trabecular subregions with identified lacunae for a NTX, ML, BIS, BIS + ML, PTH and PTH + ML sample. Boxplots for (c) trabecular bone volume fraction (BV/TV), (d) lacunar number (Lc.N) and (e) lacunar density (Lc.N/BV) between groups. Local lacunar morphometries are compared between groups for (f) local lacunar volume (<Lc.V>) and (g) local lacunar sphericity (<Lc.Sr>). (h) Schematic representation of osteocyte lacunar morphology derived from quantitative <Lc.V> and <Lc.Sr> changes across groups. Statistical significance was assessed using one-way ANOVA with Tukey’s post-hoc test for pairwise comparison. Significant differences at p < 0.05 are depicted with a symbol in the color of the corresponding group (o: to monotreatment, •: to combined treatment or sham group).

On the level of the lacunar morphology, the local lacunar volume (<Lc.V>) did not differ between the SHM- and NTX-group (Table 1, Fig. 3f). PTH treated groups however showed significantly higher <Lc.V> than NTX-, ML-, BIS and BIS + ML-groups. On the other hand, the local lacunar sphericity (<Lc.Sr>) significantly decreased for the PTH treated groups compared to all other groups (Table 1, Fig. 3g). Furthermore, the significantly increased lacunar sphericity in the NTX-group could only be reversed by ML, BIS + ML, PTH and PTH + ML treatment. Quantitatively, BIS alone did not result in a significant drop in lacunar sphericity, but it showed a visual trend of lower sphericity than the NTX-group. The global distribution of lacunar volume and sphericity further supported the noticed shift of lacunar volume towards higher values and lacunar sphericity to lower values in PTH treated groups. Notably, there are no significant differences between mono- and combined treatment for Lc.N/BV, <Lc.V> or <Lc.Sr>. The quantitative findings of <Lc.V> in Fig. 3f were qualitatively supported by the 3D visualizations in Fig. 3b. The changes in lacunar volume and sphericity by ovariectomy and treatment are illustrated in Fig. 3h. Specifically, decreased volume with increased or decreased sphericity suggests perilacunar matrix deposition and mineralization at the apices or at the sides. Conversely, concurrent increased lacunar volume and decreased sphericity may reflect osteocytic osteolysis at the apices.

### Spatial mapping reveals subregional heterogeneity in osteocyte lacunar density and morphology

By utilizing the 3D-3D multiscale registration workflow, spatial mapping of bone formation and resorption regions was performed using time-lapsed *in vivo* micro-CT mapped onto ultra-HR micro-CT to enable spatial analysis of lacunar characteristics. Four subregions were analyzed: newly formed bone (F), immediately adjacent bone (F1), middle quiescent bone (Q) and resorption-adjacent bone (R1) (Fig. 4a). Across F1, Q and R1 regions, the global treatment-response pattern of lacunar density (Lc.N/BV) (Fig. 4c) was largely preserved: the SHM-group mostly maintained significantly higher lacunar density compared to the osteoporotic groups (Table 2, Fig. 4b). Within the newly formed bone (F), however, Lc.N/BV was significantly elevated in the PTH-treated groups compared to all other groups, which exhibited lower, equivalent density values. Across all regions, no differences in group-specific trends were observed for Lc.V (Fig. 4c); specifically, the primary trend observed at the full trabecular level (Fig. 4d) was preserved within each of the four regions, with the exception of the quiescent region (Q), where Lc.V was significantly lower in the ML compared to other treatments. A similar pattern emerged for Lc.Sr. While the global treatment trends for Lc.Sr (Fig. 4e) were maintained in the F, F1 and R1 regions individually, the quiescent region showed no statistically significant differences in Lc.Sr among the SHM, NTX, ML, BIS and BIS + ML groups. Instead, Lc.Sr was significantly higher in the NTX group within the F, F1 and R1 regions, whereas PTH-treated samples consistently exhibited significantly lower Lc.Sr across all four regions compared to the other experimental groups.

**Figure 4:**
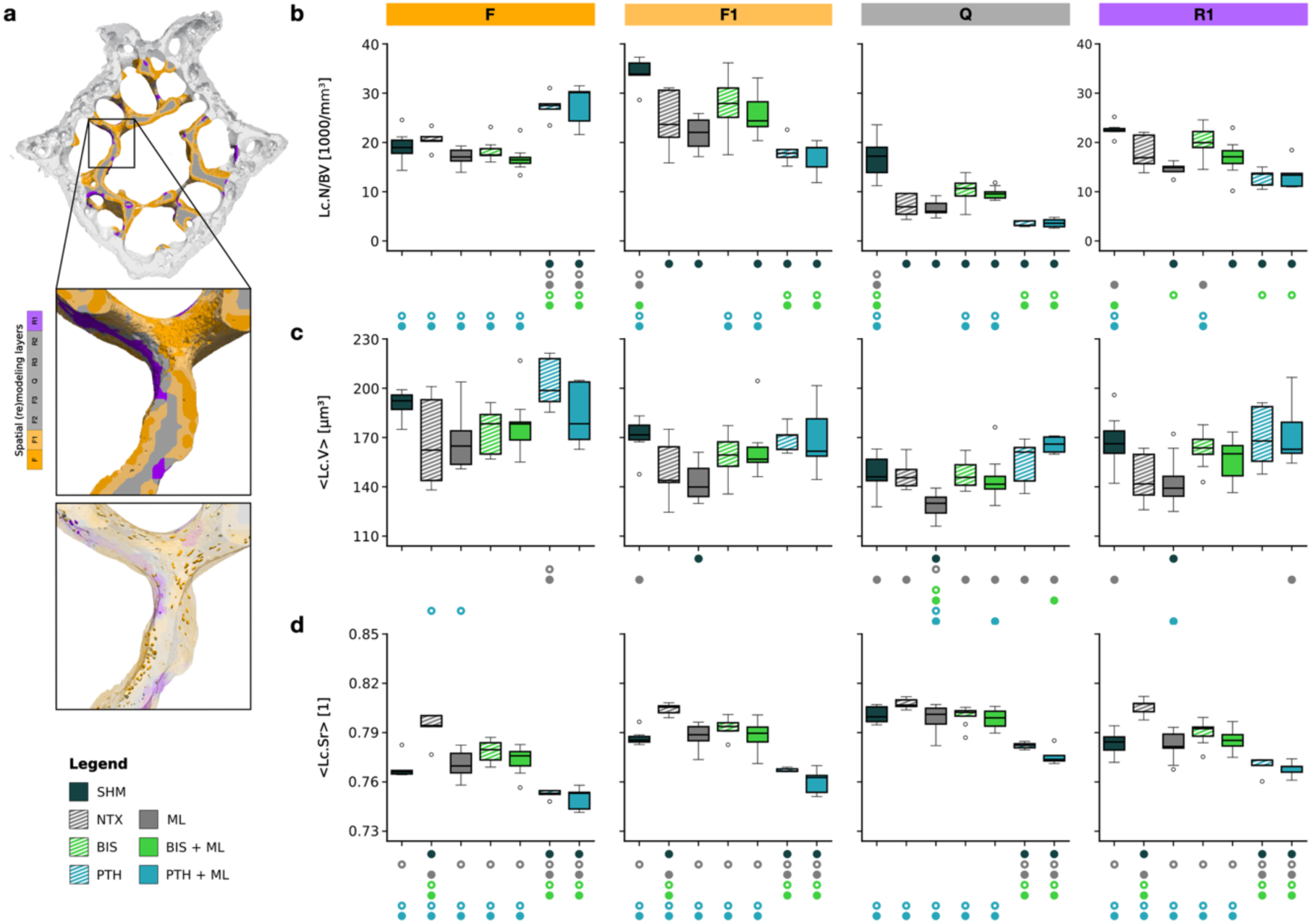
Lacunar distribution and morphometry at specific (re)modeling sites in trabecular bone compared between groups. (a) Visual representation of osteocyte lacunae analyzed in four spatial (re)modeling regions: formation (F), formation adjacent (F1), quiescent (Q) and resorption adjacent (R1). b-d) Boxplots for region-specific group differences for (b) lacunar density (Lc.N/BV), (c) lacunar volume (<Lc.V>) and (d) lacunar sphericity (Lc.Sr). Statistical significance was assessed using one-way ANOVA with Tukey’s post-hoc test for pairwise comparison. Significant differences at p < 0.05 are depicted with a symbol in the color of the corresponding group (o: to monotreatment, •: to combined treatment or sham group).

**Table 2:** Mean, standard deviation (SD) and coefficient of variation (CV) for lacunar distribution and morphology (Lc.N/BV, <Lc.V> and <Lc.Sr>) across groups and (re)modeling regions.

|  |  | Lc.N/BV |  | <Lc.V> |  | <Lc.Sr> |  |
| --- | --- | --- | --- | --- | --- | --- | --- |
| | | Mean ± SD<br>[1000/$\mu\text{m}^3$] | CV<br>[%] | Mean ± SD<br>[$\mu\text{m}^3$] | CV<br>[%] | Mean ± SD<br>[$\mu\text{m}^3$] | CV<br>[%] |
| **SHM** | **F** | 19.191 ± 3.185 | 16.596 | 190.320 ± 8.469 | 4.450 | 0.768 ± 0.006 | 0.838 |
| | **1** | 34.198 ± 2.864 | 8.376 | 170.651 ± 11.611 | 6.804 | 0.787 ± 0.005 | 0.590 |
| | **Q** | 16.829 ± 4.302 | 25.561 | 148.257 ± 12.206 | 8.233 | 0.801 ± 0.005 | 0.652 |
| | **-1** | 22.572 ± 1.463 | 6.481 | 167.576 ± 16.968 | 10.126 | 0.783 ± 0.007 | 0.937 |
| **NTX** | **F** | 20.524 ± 2.133 | 10.394 | 167.641 ± 28.374 | 16.925 | 0.793 ± 0.010 | 1.226 |
| | **1** | 24.438 ± 6.475 | 26.494 | 150.086 ± 19.823 | 13.208 | 0.804 ± 0.004 | 0.449 |
| | **Q** | 7.198 ± 2.411 | 33.489 | 147.549 ± 9.679 | 6.560 | 0.808 ± 0.003 | 0.400 |
| | **-1** | 17.992 ± 3.620 | 20.121 | 145.215 ± 16.031 | 11.040 | 0.806 ± 0.005 | 0.680 |
| **ML** | **F** | 17.237 ± 1.695 | 9.831 | 169.783 ± 18.953 | 11.163 | 0.771 ± 0.008 | 0.993 |
| | **1** | 21.725 ± 3.014 | 13.873 | 143.771 ± 11.782 | 8.195 | 0.788 ± 0.007 | 0.901 |
| | **Q** | 6.546 ± 1.488 | 22.736 | 128.189 ± 8.125 | 6.338 | 0.799 ± 0.008 | 0.979 |
| | **-1** | 14.669 ± 1.145 | 7.807 | 143.335 ± 15.524 | 10.831 | 0.783 ± 0.009 | 1.152 |
| **BIS** | **F** | 18.323 ± 2.086 | 11.385 | 173.440 ± 13.700 | 7.899 | 0.779 ± 0.006 | 0.805 |
| | **1** | 27.514 ± 6.101 | 22.175 | 158.522 ± 12.544 | 7.913 | 0.793 ± 0.005 | 0.635 |
| | **Q** | 9.969 ± 2.869 | 28.779 | 147.448 ± 8.446 | 5.728 | 0.800 ± 0.006 | 0.736 |
| | **-1** | 20.098 ± 3.333 | 16.582 | 162.681 ± 10.609 | 6.521 | 0.790 ± 0.007 | 0.893 |
| **BIS + ML** | **F** | 16.722 ± 2.503 | 14.966 | 178.067 ± 17.193 | 9.655 | 0.774 ± 0.009 | 1.103 |
| | **1** | 25.789 ± 3.949 | 15.314 | 161.594 ± 17.441 | 10.793 | 0.788 ± 0.009 | 1.147 |
| | **Q** | 9.593 ± 1.246 | 12.989 | 144.837 ± 13.807 | 9.533 | 0.798 ± 0.006 | 0.723 |
| | **-1** | 16.899 ± 3.520 | 20.831 | 155.846 ± 13.417 | 8.609 | 0.785 ± 0.007 | 0.872 |
| **PTH** | **F** | 27.310 ± 2.709 | 9.918 | 203.037 ± 15.899 | 7.831 | 0.753 ± 0.003 | 0.364 |
| | **1** | 18.226 ± 2.741 | 15.039 | 169.484 ± 8.280 | 4.885 | 0.767 ± 0.001 | 0.154 |
| | **Q** | 3.472 ± 0.555 | 15.992 | 154.716 ± 14.212 | 9.186 | 0.782 ± 0.002 | 0.258 |
| | **-1** | 12.838 ± 1.857 | 14.467 | 170.205 ± 19.396 | 11.396 | 0.770 ± 0.006 | 0.734 |
| **PTH + ML** | **F** | 27.574 ± 4.335 | 15.722 | 183.713 ± 19.570 | 10.653 | 0.750 ± 0.007 | 0.946 |
| | **1** | 17.026 ± 3.514 | 20.636 | 169.550 ± 22.239 | 13.116 | 0.760 ± 0.008 | 1.021 |
| | **Q** | 3.637 ± 0.921 | 25.326 | 165.660 ± 5.094 | 3.075 | 0.776 ± 0.006 | 0.712 |
| | **-1** | 13.517 ± 3.028 | 22.402 | 172.706 ± 21.059 | 12.193 | 0.768 ± 0.005 | 0.627 |

Abbreviations: SHM: sham group; NTX: untreated ovariectomized control group; ML: mechanically loaded group; BIS: bisphosphonate treated group; BIS+ML: bisphosphonate treated and mechanically loaded group; PTH: parathyroid hormone treated group; PTH+ML: parathyroid hormone treated and mechanically loaded group; F: formation region, 1: formation adjacent region 1, Q: quiescent region; -1: resorption adjacent region -1; Lc.N/BV: lacunar density; <Lc.V>: local lacunar volume; <Lc.Sr>: local lacunar sphericity. SD: standard deviation, CV: coefficient of variation

Beyond inter-group comparisons, isolating the spatial profile of the lacunar morphometric parameters within the PTH + ML-treated group highlighted an intra-individual gradient across functional bone regions (Fig. 5). PTH + ML-treated samples demonstrated a distinct spatial transition: lacunar expansion and elongation were most pronounced in newly formed bone (F) and dampened in quiescent bone (Q). Visualizing these parameter distributions in 3D indicated that PTH + ML-induced lacunar remodeling is not uniformly distributed throughout the trabecular compartment but is spatially concentrated in regions of active tissue turnover. All intra-individual boxplots of Lc.N/BV, Lc.V and Lc.Sr are provided in Supplementary Figure 5.

**Figure 5:**
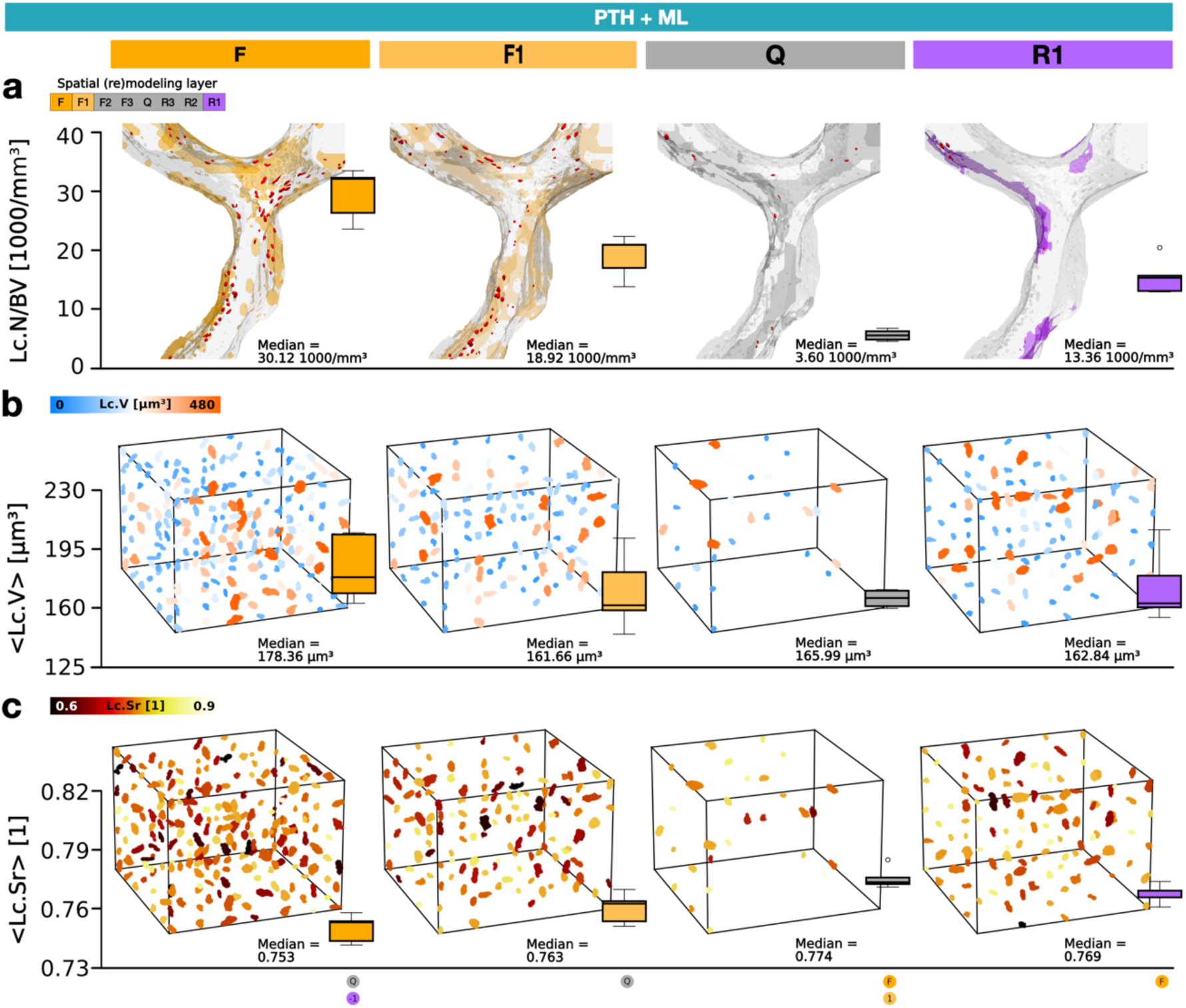
Lacunar distribution and morphometry of the PTH + ML compared between (re)modeling sites. Boxplots for (a) lacunar density (Lc.N/BV), (b) local lacunar volume (Lc.V) and (c) local lacunar sphericity (Lc.Sr) for PTH + ML in four spatial (re)modeling regions: formation (F), formation adjacent (F1), quiescent (Q) and resorption adjacent (R1). b-c) Representative volume elements (RVE, volume = 0.008 µm^3^) are included alongside the boxplots to visually model the data, where lacunar density is reflected by the number of lacunae within each RVE, and the colormaps indicate Lc.V and Lc.Sr. To populate the RVEs, lacunae within one standard deviation of the mean regional Lc.V were randomly sampled until the targeted group density was achieved.

## 3. Discussion

By establishing a 3D-3D multiscale registration framework, this study reveals that physical and pharmacological osteoporosis therapies differentially modify the osteocyte lacunar network through distinct cell-scale perilacunar adaptations. Integrating time-lapsed *in vivo* micro-CT with *ex vivo* ultra-HR micro-CT enabled the direct mapping of endpoint 3D osteocyte lacunar distribution and morphology onto the bone’s dynamic micro-CT derived (re)modeling history. The spatial integration moves beyond whole-bone averages, demonstrating that both osteoporotic degeneration and therapeutic osteocyte lacunar remodeling are highly heterogenous across functional microenvironments. At the cellular scale, systemic anabolic PTH acted as a dominant driver of osteocytic perilacunar remodeling (PLR), including lacunar expansion and elongation that reached a morphometric saturation point, superseding any synergistic effect from concurrent mechanical loading (PTH + ML). Meanwhile, bisphosphonates stabilized the lacunar architecture at levels comparable to healthy bone, with mechanical stimulation conferring no additional morphological benefit when combined with BIS treatment (BIS + ML). Crucially, spatial analysis across local bone regions revealed that estrogen loss disrupts the natural maturation gradient of lacunar sphericity from surface-adjacent (re)modeling sites to the interior matrix core, whereas the investigated therapies selectively counteract this disruption or systematically shift the overall lacunar morphometric pattern. Together, these morphometric and spatial signatures demonstrate that osteocytes actively tune their local microenvironment through perilacunar adaptations in response to physical, endocrine and matrix-targeted cues. These findings support our primary hypotheses, confirming that while bisphosphonates stabilize the local lacunar architecture, PTH triggers perilacunar adaptation over the whole investigated trabecular bone.

Osteocytes secrete regulatory factors that not only orchestrate surface osteoblasts and osteoclast activity, but also act locally to regulate PLR^9^ to actively tune the degree to which they sense strain.^34^ In our model, ovariectomy induced modest alterations in lacunar volume alongside a marked increase in lacunar sphericity, adding to previous evidence of PLR where estrogen deficiency disrupts perilacunar matrix architecture primarily seen through canalicular expansion.^35^ Conversely, PTH treatment significantly expanded lacunar volume while decreasing sphericity, reflecting targeted pericellular matrix resorption and architectural adaptation. These morphometric shifts corroborate earlier findings demonstrating that systemic PTH signaling directly activates osteocytic matrix degradation through enzymes such as cathepsin K and matrix metalloproteinases.^36,37^ Biologically, these reciprocal morphometric shifts point to directional perilacunar remodeling, where localized osteocytic osteolysis and perilacunar matrix deposition and mineralization selectively reshape the void geometry. The reduction in sphericity alongside expanded lacunar volume under systemic PTH treatment suggests that osteocytic osteolysis occurs asymmetrically.

A central finding of our lacunar morphometric analysis in different (re)modeling regions is how pharmacological and mechanical interventions modulate the spatial gradient of lacunar sphericity. *In vitro* evidence suggests that osteocyte geometry directly dictates their mechanosensitivity, with elongated, less spherical cells exhibiting lower mechanosensitivity compared to rounder osteocytes.^17^ In healthy controls, mechanically loaded, BIS-treated and PTH-treated groups, lacunar sphericity followed a distinct developmental gradient corresponding to functional (re)modeling microenvironments: sphericity was lowest in newly formed bone, intermediate in formation-and resorption-close regions, and reached its highest level in mature, quiescent bone. This gradient reflects the natural morphological maturation of osteocytes as newly entrapped cells transition from an elongated phenotype toward mature lacunar architecture. Biologically, the lower sphericity observed in newly formed bone could act as a regulatory mechanism for maintaining lower inherent cellular mechanosensitivity in young matrix to prevent excessive bone formation response to physiological strains during active mineralization. Estrogen deficiency through ovariectomy disrupted this spatial maturation. Non-treated ovariectomized samples exhibited a uniformly elevated sphericity across all spatial domains. This early rounder shape of osteocyte lacunae within newly formed bone suggests that estrogen loss induces a state of pathological hyper-mechanosensitivity in the young matrix, driving premature pericellular degradation. Both mechanical loading and BIS treatment successfully prevented this early potentially pathological rounding, maintaining significantly lower sphericity in newly formed bone. Systemic PTH treatment also preserved this development gradient but shifted the spatial profile of lacunar sphericity downward across all investigated bone regions.

Because systemic PTH acts on osteocytes across the entire bone tissue, it drives perilacunar remodeling (PLR) throughout the investigated trabecular bone towards an elongated, lower-sphericity phenotype across all spatial regions. Mirroring this trend, lacunar volume followed an inverse gradient with values peaking in newly formed bone and lowest values in quiescent bone further supporting a directional PLR over all microenvironments for all groups except the NTX-group.

In addition to the cell-scale morphological adaptations, spatial mapping of two micro-CT derived data volumes revealed distinct distribution patterns in response to the investigated treatments in osteocyte lacunar density. Crucially, while both physical and pharmacological interventions successfully restored tissue-level bone volume fraction, global trabecular lacunar density remained lower relative to healthy controls. Reduced osteocyte density in osteoporotic human samples has been reported previously^38,39^ but opposite observations have also been reported.^21,40,41^ Qiu et al.^42^ reported an inverse relationship of bone formation rate and osteocyte density as a sort of negative feedback system. Increased lacunar density and thus presumed osteocyte density in newly formed bone in PTH treated groups could therefore be an additional restraining factor to prohibit excessive bone formation.

Our cellular-level findings provide a direct basis for recent tissue-level observations by Schulte et al. regarding treatment synergies and mechanotransduction.^33^ Consistent with their macroscale findings and validated by our static and dynamic bone morphometry, BIS preserved trabecular number but yielded no additive anabolic effect when combined with mechanical loading, suggesting a diminished osteoblast response to mechanical signals. At the cellular scale, our spatial (re)modeling analysis of lacunar morphometry shows that both BIS and ML independently enforce elongated, lower-sphericity phenotype in newly formed bone. BIS maintains osteocytes in a less mechanosensitive state within young matrix, capping further strain amplification and providing a precise cellular mechanism for the lack of additive tissue-level responses under combined BIS and ML, directly supporting our hypothesis of a stabilized, morphologically unreactive cellular network. Conversely, at the tissue level, combining anabolic PTH with mechanical loading drives additive structural changes. Notably, while Schulte et al. demonstrated this macroscale additive effects using a 40 µg/kg PTH treatment regimen, our tissue-level data confirms that this additive effect persists even at a higher dose of 80 µg/kg. At the single-cell scale, our data reveals that while this higher PTH dose drives lacunar expansion and elongation that reaches a morphometric saturation, it maintains high lacunar density and active pericellular turnover. Refining our original hypothesis which predicted continuous dynamic lacunar morphological adaptation, we demonstrate that even when individual lacunar geometry reaches a physical remodeling ceiling, this preserved network viability allows the osteocyte population to continually translate mechanical strain into bone formation. Furthermore, Schulte et al. observed that despite minimal overall bone turnover, BIS-treated tissue efficiently targeted mechanical disuse areas, hypothesizing a compensatory osteocytic mechanosensitivity. Our spatial mapping supports this model: across all groups, high lacunar density is naturally retained in newly formed and formation-close regions compared to quiescent older bone. In BIS-treated bone, this dense structurally intact osteocyte network supports the theory of concentrated localized disuse signaling (such as apoptotic RANKL release)^43^ in low strain energy density (SED) regions, enabling targeted osteoclast recruitment to mechanically redundant sites despite suppressed global turnover.

Several methodological and biological limitations should be noted. While dynamic (re)modeling regions derived from *in vivo* micro-CT represent localized matrix microenvironments rather than single-cell pericellular boundaries, the robust subregion-specific morphometric gradients observed across functional sites demonstrate that this multiscale mapping captures distinct cellular adaptation zones. Furthermore, the ultra-HR micro-CT measures mineralized lacunar voids rather than direct soft-tissue osteocyte cellular morphology, cytoskeletal dynamics or fluid flow in the LCN. Thus, 3D lacunar volume and sphericity serve as morphological approximations for PLR, representing a cumulative structural outcome of past mechanical and cellular activity rather real-time functional signaling. Additionally, due to ultra-HR field of view and computational processing constraints, cell-scale lacunar analysis was conducted within a standardized sub-volume of the trabecular compartment rather than the entire vertebra. However, this sub-region was selected to be representative of the overall tissue structure, as evidenced by BV/TV changes in the HR domain closely mirroring the whole-trabecular BV/TV endpoint values observed *in vivo*. In terms of biological interpretation, standard X-ray micro-CT cannot distinguish between occupied and empty lacunae. Finally, while our time-lapsed *in vivo* micro-CT image protocol captured distinct functional (re)modeling phases, future studies utilizing multi-time-point nanoscale imaging could not only track the individual lacunae and osteocytes but also add valuable morphological knowledge on the canalicular system connecting individual osteocytes.

In summary, by establishing a 3D-3D multiscale registration framework, this study bridges the gap between dynamic tissue-level (re)modeling and cell-scale perilacunar morphological adaptations. We demonstrate that physical and pharmacological interventions differentially alter the osteocyte lacunae. The new insights of changes in lacunar distribution and morphology will hopefully contribute to a better understanding of the mechanisms underlying disease and treatment in postmenopausal osteoporosis.

## 4. Methods

### Experimental design

Our objective was to investigate spatial differences in osteocyte lacunar morphology across healthy, osteoporotic and treated osteoporotic bone. The study design is illustrated in Fig. 1a. Out of 50 15-week-old C57BL/6 female mice (C57BL/JRccHsd, RCC/Harlan; Füllinsdorf, Switzerland/Horst, The Netherlands), 43 were subjected to bilateral ovariectomy to induce estrogen depletion and establish a mouse model of osteoporosis. The other 7 mice underwent sham surgery without ovariectomy (SHM-group). The ovariectomized mice were divided into 6 groups and received their respective treatment after 5 weeks of recovery: untreated ovariectomized controls (NTX, n = 5), ovariectomized controls with ML (ML, n = 9), BIS (n = 9), BIS + ML (n = 10), PTH (n = 5), and PTH + ML (n = 5). The animals in the BIS and BIS + ML groups were reused from the cohort previously published by Schulte et al.^33^ To allow direct comparison with the new groups (SHM, NTX, ML, PTH and PTH + ML), previously reported static and dynamic *in vivo* morphometric parameters are included here, whereas ultra-HR derived lacunar parameters represent new analyses of these samples. One mouse in the BIS + ML was excluded from analysis as it did not lose bone volume after ovariectomy, resulting in 9 animals in this group. BIS (100 µm/kg zoledronate, Zometa 4 mg/5 ml; Novartis Pharma Schweiz AG, Bern) was administered once at week 20 of age while PTH (80 µg/kg PTH, hPTH 1-34, Bachem AG, Switzerland) was administered daily over four weeks. Mechanical loading (ML) was applied to the 6th caudal vertebra through a highly controlled cyclic mechanical loading model (8 N, 10 Hz, 3000 cycles).^44^ The loading protocol has previously been shown to cause a 20% increase in the trabecular bone volume fraction (BV/TV) of healthy mice.^45^ *In vivo* micro-CT was performed at weeks 15, 20, 22 and 24. At the end of the study, the vertebra was extracted, embedded in poly(methyl methacrylate) (PMMA) and a high-resolution *ex vivo* micro-CT was performed. While the data was gained in three experiments at different times, the study and loading parameters were the same except the ovariectomy and administered treatment.

### Ethics statement

All mouse experiments were performed in accordance with relevant national regulations (Swiss Animal Welfare Act, TSchG, and Swiss Animal Welfare Ordinance, TSchV) and authorized by the Zürich Cantonal Veterinary Office (approved license number: 171/2008); Kantonales Veterinäramt Zürich, Switzerland)

### *In vivo* micro-CT: static and dynamic morphometry

*In vivo* micro-CT images were obtained of the 6^th^ caudal vertebra with a vivaCT 40 (Scanco Medical AG, Brüttisellen, Switzerland) at a nominal low resolution of 10.5 µm as previously detailed.^33^ Follow up scans were registered to the previous scan, a constrained 3D gaussian filter (σ = 1.2, support = 1) was applied followed by a thresholding procedure (threshold at 22%, approximately 580 mg HA/cm^3^). Automated trabecular masks were created^32^ and trabecular static bone morphometry was calculated for each scan according to Bouxsein et al.^46^ Dynamic bone morphometry was performed using the registered scans and superimposing segmented bone volumes of consecutive time points.^47^

### Ultra-high resolution (ultra-HR) micro-CT: osteocyte lacunar morphology

Following surgical extraction, the 6^th^ caudal vertebra was dehydrated in graded ethanol, cleared in xylene and embedded in poly(methyl methacrylate) (PMMA). A 1.1 mm thick volume along the longitudinal axis of the vertebra was then imaged using a µCT50 (Scanco Medical AG, Brüttisellen, Switzerland) at high nominal resolution of 1.2 µm. The scanner was operated with a 0.5 mm aluminum filter, 57 µA current, 4 W power, 70 kVp energy, 1.5 s integration time, level 6 data averaging and with a total of 1500 projections per 180°. Before each high-resolution scan, a pre-scan (1 h) was performed to warm the sample and reduce motion artefacts caused by thermal effects. A constrained 3D gaussian filter (σ = 0.8, support = 1) followed by an individualized threshold based on the tissue mineral density distribution of each sample was applied to segment osteocyte lacunae^48^. Only void volumes in the range of 50-2000 µm^3^ were accepted for analysis to reflect the range of osteocyte lacunar volumes.^49^ Lacunar morphometric parameters were calculated using a custom validated Python script (v1.10.4, Python Software Foundation, Delaware, USA) in XamFlow (Lucid AG, Zurich, Switzerland).^48^ The following parameters were evaluated in the analysis: Lacunar number (Lc.N), lacunar density (Lc.N/BV) and lacunar volume (Lc.V) as first defined by Mader et al.^50^ and lacunar sphericity (Lc.Sr) as defined by Akhter et al.^51^ Bone volume in this case was calculated as voxel volume. Lc.V and Lc.Sr were reported as local lacunar parameters normalized to the number of lacunae in one sample (and region) and denoted with angle brackets (< >). Group-level medians were calculated from the individual sample means. Group variability was quantified using the coefficient of variation (CV), calculated as the group SD divided by the group mean.

### 3D-3D multiscale registration workflow

To investigate how osteocyte lacunar distribution and morphology varies across local bone (re)modeling microenvironments, the time-lapsed *in vivo* micro-CT images from week 22 and 24 and the respective (re)modeling sides were aligned with the corresponding high-resolution micro-CT image. The detailed workflow is visualized in Fig. 1b and further explained in the following sections.

#### Ex vivo micro-CT image rescaling

Ultra high-resolution (ultra-HR) micro-CT images were rescaled to 10.5 µm nominal resolution (cubic b-spline interpolation) using a scikit-image implementation (Fig. 1b(1, 2)).^52^ The following steps of 3D-3D grayscale registration of the *in vivo* and high-resolution micro-CT and the subsequent steps of (re)modeling site extraction, surface constrained projection to the ultra-HR micro-CT and distance mapping were performed at the lower nominal resolution of 10.5 µm to reduce computational demands. Micro-CT image registration at this resolution has also been demonstrated successfully in previous studies.^47^ Raw ultra-HR micro-CT images were cropped prior to processing to avoid dimension inconsistencies introduced by resampling during downscaling and subsequent upscaling. This ensured that the image dimensions remained constant and could be restored without rounding-related mismatches after interpolation onto coarser or finer voxel grids and associated discretization effects. A nominal scaling factor of 8.750 (from 1.2 µm to 10.5 µm voxel size) achieved a one-to-one voxel match between the *in vivo* and ultra-HR image after downscaling but did not result in an optimal match of the scanned vertebra due to minor tissue deformations introduced during sample embedding, as well as differences in scanning geometry and reconstruction between the *in vivo* and *ex vivo* images (Supplementary Figure 3a). Therefore, the scaling factor was adjusted to an effective scaling factor of 8.569. It was calculated as the mean of the minimal average symmetric surface distance (ASSD) between the registered binarized *in vivo* and HR scans (threshold at 22%, approximately 580 mg HA/cm^3^) for each group (Supplementary Figure 3). The effective voxel size of the downscaled ultra-HR micro-CT resulted in 10.3 µm, 2% smaller than the reported 10.5 µm by the micro-CT system.

#### 3D-3D grayscale registration

The 1.1 mm thick volume of the HR micro-CT was visually identified on the with each other registered w22 and w24 *in vivo* micro-CT images, and the corresponding volume of interest (VOI) was cropped in both images. Additional padding of 20 voxels in z-direction at both ends was applied to the *in vivo* and downscaled ultra-HR images to compensate for potential information loss due to transformations during the following registration. All images were 3D Gaussian filtered (σ = 1.2, support = 1) prior to the grayscale-based registration. The w24 *in vivo* image was registered to the downscaled ultra-HR image and the resulting transformation matrix applied to the *in vivo* registered w22 image (Fig. 1b(3)) and thus effectively matching up both *in vivo* and the HR images. A global thresholding of 22% (approximately 580 mg HA/cm^3^) was applied to the *in vivo* micro-CT images before superimposing the segmented bone volumes to determine (re)modeling sites within the cropped VOI (Fig. 1b(4, 6)). The *in vivo* (re)modeling sites were then superimposed on the downscaled ultra-HR image (Fig. 1b(5, 7)).

#### Surface constrained projection, distance mapping and upscaling

To align the (re)modeling sites with the exact bone boundaries of downscaled ultra-HR image, the 3D binary bone volume of the w24 scan was transformed to the bone volume of the HR scan using a SciPy implementation of a Euclidean distance transform.^53^ Thresholding the interpolated distance transform resulted in intermediate binary volumes. The labeled (re)modeling data was then stepwise transferred to the nearest bone surface voxel in the evolving shape (4 steps). Within each step, formation volumes were forced to fall to the next voxel within the bone boundaries whereas resorption volumes were assigned outside the bone. This ensured correct spatial assignment of formation and resorption volumes to the ultra-HR bone surface at the end of the transformation (Fig. 1b(8)). Following the surface constrained projection, the individual 3D formation and resorption distance maps on the downscaled ultra-HR micro-CT were created using the same Euclidean distance transform implementation of SciPy.^53^ The regions were cut off at three surrounding layers (1 to 3 for formation, -1 to -3 for resorption) and remaining voxels were defined as purely quiescent regions (Q) (Fig. 1b(9, 10)). The individual spatial (re)modeling maps were upscaled to 1.2 µm nominal resolution using trilinear interpolation (Fig. 1b(11, 12)). The resampled values were discretized using an offset-based rounding scheme (x + 0.3, rounded to the nearest integer) to ensure volume conservation in the (re)modeling regions. The commonly used nearest-neighbor interpolation was avoided to prevent jagged edges in the upscaled spatial (re)modeling map. The final spatial (re)modeling map was derived by voxel-wise integration of the formation and resorption distance maps (Fig. 1b(13)), reducing the bivariate spatial map into eight distinct (re)modeling classes for each voxel: F, F1, F2, F3, Q, R3, R2, R1. Note that the resorption volume could not be analyzed in the HR image as it is a non-existent volume on the scan. While it is possible to keep multiple distance layers for analysis, subsequent analyses were restricted to the formation region, the immediately adjacent formation layer F1 and the immediate adjacent resorption layer R1. All other middle regions were summarized to a combined quiescent region (Q). This simplification reduced analytical complexity while preserving the spatial regions most directly associated with active bone formation and resorption. For each sample, the lacunar parameter means and standard deviation (SD) was calculated for the above-mentioned regions. Group-level medians were calculated from the sample means.

#### Quantitative Assessment of the Workflow

To justify the accuracy of the 3D-3D multiscale registration workflow in the absence of a ground truth, it was assessed at key image processing steps through quantitative evaluation of established, standardized dynamic bone morphometric parameters: bone formation rate (BFR), bone resorption rate (BRR), mineral apposition rate (MAR), mineral resorption rate (MRR), mineralizing surface (MS) and eroded surface (ES) (Supplementary Material 1.1, Supplementary Figure 1). The calculation of the dynamic parameters was implemented in XamFlow (Lucid AG, Zürich, Switzerland) and validated against the existing pipeline in IPL (Supplementary Figure 2).^47^

### Statistical analysis

One-way ANOVA was used to test all differences within and between groups and regions. Before assessing statistical significance between groups, usually the data is being tested on normality and homogeneity of variance. The groups tested in this study have an n = 5-9 per group which rendered normality tests underpowered. Normality in our datasets was thus only assumed based on evidence from previous data on static and dynamic morphometry^33^ and large-scale assessment of local osteocyte lacunar parameters.^54^ If the ANOVA was significant, Tukey’s HSD post-hoc test was run for all comparisons. All tests were performed in Python (version 3.12.12) using the open-source libraries SciPy^53^ (version 1.16.1) and statsmodels (version 0.14.5).^55^ If not indicated otherwise, significant differences at p < 0.05 are depicted with a symbol in the color of the corresponding group (o: to monotreatment, • : to combined treatment or sham group, see Supplementary Table 2 for color encoding).

## Data availability

All data are available in the main text and Supplementary Information.

## Supplementary information

Supplementary information accompanies the manuscript on the *Bone Research* website http://www.nature.com/boneres.

## Acknowledgments

Funding was provided by the Swiss National Science Foundation (Sinergia Grant 213520; SL) and the European Union grant VPHOP (FP7-ICT-2008-223865; CW, FML, FAS, GAK). The authors would like to thank Lucid AG for assisting with the 3D-3D multiscale registration workflow evaluation study and Carmen Eskesen for early contribution to the project. During the preparation of this work, the authors used OpenAI ChatGPT and Gemini to guide and improve the readability of this manuscript and assist in writing custom data analysis scripts. After using these tools, the authors reviewed, tested and edited the text and code as needed and take full responsibility for the content of the publication.

## Contributions

1. Designed the work that led to the submission, acquired data, and/or played an important role in interpreting the results
2. Drafted or revised the manuscript
3. Approved the final version

Sara Lindenmannn (1, 2, 3), Zihui Li (1, 3) Claudia Weigt (1, 3), Floor M. Lambers (1, 3), Friederike A. Schulte (1, 2, 3), Gisela A. Kuhn (1, 2, 3), Neashan Mathavan (1, 2, 3), Ralph Müller (1, 2, 3)

## References

1. Wolff, J. Das Gesetz der Transformation der Knochen. (Schattauer, Stuttgart, 1991).

2. Roux, W. Der Kampf Der Theile Im Organismus. Ein Beitrag Zur Vervollständigung Der Mechanischen Zweckmässigkeitslehre, von Wilhelm Roux. (W. Engelmann, Leipzig, 1881). doi:10.5962/bhl.title.1491.

3. Mullender, M. G., Huiskes, R., Versleyen, H. & Buma, P. Osteocyte density and histomorphometric parameters in cancellous bone of the proximal femur in five mammalian species. J. Orthop. Res. 14, 972–979 (1996).

4. Bonewald, L. F. The amazing osteocyte. J. Bone Miner. Res. 26, 229–238 (2011).

5. Hillsley, M. V. & Frangos, J. A. Review: Bone tissue engineering: The role of interstitial fluid flow. Biotechnol. Bioeng. 43, 573–581 (1994).

6. Weinbaum, S., Cowin, S. C. & Zeng, Y. A model for the excitation of osteocytes by mechanical loading-induced bone fluid shear stresses. J. Biomech. 27, 339–360 (1994).

7. Duncan, R. L. & Turner, C. H. Mechanotransduction and the functional response of bone to mechanical strain. Calcif. Tissue Int. 57, 344–358 (1995).

8. Mcgarry, J. G., Klein-Nulend, J., Mullender, M. G. & Prendergast, P. J. A comparison of strain and fluid shear stress in stimulating bone cell responses—a computational and experimental study. FASEB J. 19, 1–22 (2005).

9. Yee, C. S., Schurman, C. A., White, C. R. & Alliston, T. Investigating Osteocytic Perilacunar/Canalicular Remodeling. Curr. Osteoporos. Rep. 17, 157–168 (2019).

10. Nicolella, D. P., Moravits, D. E., Gale, A. M., Bonewald, L. F. & Lankford, J. Osteocyte lacunae tissue strain in cortical bone. J. Biomech. 39, 1735–1743 (2006).

11. Muñoz, A., De Paolis, A., Cardoso, L. & Carriero, A. Osteocyte-lacuna shape and canaliculi architecture dictate fluid flow around osteocyte, and strain of cell and bone matrix: implications for cell mechanobiology and bone fragility. Bone 200, 117613 (2025).

12. Van Hove, R. P. et al. Osteocyte morphology in human tibiae of different bone pathologies with different bone mineral density — Is there a role for mechanosensing? Bone 45, 321–329 (2009).

13. Carriero, A. et al. Altered lacunar and vascular porosity in osteogenesis imperfecta mouse bone as revealed by synchrotron tomography contributes to bone fragility. Bone 61, 116–124 (2014).

14. Mullender, M. G., Van Der Meer, D. D., Huiskes, R. & Lips, P. Osteocyte density changes in aging and osteoporosis. Bone 18, 109–113 (1996).

15. Carter, Y., Thomas, C. D. L., Clement, J. G. & Cooper, D. M. L. Femoral osteocyte lacunar density, volume and morphology in women across the lifespan. J. Struct. Biol. 183, 519–526 (2013).

16. Gatti, V., Azoulay, E. M. & Fritton, S. P. Microstructural changes associated with osteoporosis negatively affect loading-induced fluid flow around osteocytes in cortical bone. J. Biomech. 66, 127–136 (2018).

17. Bacabac, R. G. et al. Round versus flat: Bone cell morphology, elasticity, and mechanosensing. J. Biomech. 41, 1590–1598 (2008).

18. Consensus development conference: Diagnosis, prophylaxis, and treatment of osteoporosis. Am. J. Med. 94, 646–650 (1993).

19. Tomkinson, A., Gevers, E. F., Wit, J. M., Reeve, J. & Noble, B. S. The Role of Estrogen in the Control of Rat Osteocyte Apoptosis. J. Bone Miner. Res. 13, 1243–1250 (1998).

20. Manolagas, S. C. From Estrogen-Centric to Aging and Oxidative Stress: A Revised Perspective of the Pathogenesis of Osteoporosis. Endocr. Rev. 31, 266–300 (2010).

21. Mullender, M. G., Van Der Meer, D. D., Huiskes, R. & Lips, P. Osteocyte density changes in aging and osteoporosis. Bone 18, 109–113 (1996).

22. Vashishth, D., Verborgt, O., Divine, G., Schaffler, M. B. & Fyhrie, D. P. Decline in osteocyte lacunar density in human cortical bone is associated with accumulation of microcracks with age. Bone 26, 375–380 (2000).

23. Ma, Y.-L. et al. Quantitative associations between osteocyte density and biomechanics, microcrack and microstructure in OVX rats vertebral trabeculae. J. Biomech. 41, 1324– 1332 (2008).

24. Reid, I. R. Bisphosphonates in the treatment of osteoporosis: a review of their contribution and controversies. Skeletal Radiol. 40, 1191–1196 (2011).

25. Luckman, S. P., Hughes, D. E., Coxon, F. P., Russell, R. G. G. & Dr.Rogers, M. J. Nitrogen-Containing Bisphosphonates Inhibit the Mevalonate Pathway and Prevent Post-Translational Prenylation of GTP-Binding Proteins, Including Ras. J. Bone Miner. Res. 13, 581–589 (1998).

26. Jilka, R. L. et al. Increased bone formation by prevention of osteoblast apoptosis with parathyroid hormone. J. Clin. Invest. 104, 439–446 (1999).

27. Inaoka, T. et al. Sequential Analysis of Gene Expression after an Osteogenic Stimulus - c-fos Expression Is Induced in Osteocytes. Biochem. Biophys. Res. Commun. 217, 264–270 (1995).

28. Kleinnulend, J., Semeins, C. M., Ajubi, N. E., Nijweide, P. J. & Burger, E. H. Pulsating Fluid Flow Increases Nitric Oxide (NO) Synthesis by Osteocytes but Not Periosteal Fibroblasts - Correlation with Prostaglandin Upregulation. Biochem. Biophys. Res. Commun. 217, 640–648 (1995).

29. Forwood, M. R. Inducible cyclo-oxygenase (COX-2) mediates the induction of bone formation by mechanical loading in vivo. J. Bone Miner. Res. 11, 1688–1693 (1996).

30. Ajubi, N. E., Klein-Nulend, J., Alblas, M. J., Burger, E. H. & Nijweide, P. J. Signal transduction pathways involved in fluid flow-induced PGE_2_ production by cultured osteocytes. Am. J. Physiol.-Endocrinol. Metab. 276, E171–E178 (1999).

31. Schulte, F. A. et al. Combined physical and pharmacological anabolic osteoporosis therapies increase bone response and mechanoregulation in female mice. Nat. Commun. 17, 3759 (2026).

32. Lambers, F. M., Schulte, F. A., Kuhn, G., Webster, D. J. & Müller, R. Mouse tail vertebrae adapt to cyclic mechanical loading by increasing bone formation rate and decreasing bone resorption rate as shown by time-lapsed in vivo imaging of dynamic bone morphometry. Bone 49, 1340–1350 (2011).

33. Schulte, F. A. et al. Combined physical and pharmacological anabolic osteoporosis therapies increase bone response and mechanoregulation in female mice. Nat. Commun. 17, 3759 (2026).

34. Rubin, C., Sun, Y., Hadjiargyrou, M. & McLeod, K. Increased expression of matrix metalloproteinase-1 in osteocytes precedes bone resorption as stimulated by disuse: Evidence for autoregulation of the cell’s mechanical environment? J. Orthop. Res. 17, 354–361 (1999).

35. Sharma, D. et al. The effects of estrogen deficiency on cortical bone microporosity and mineralization. Bone 110, 1–10 (2018).

36. Wysolmerski, J. J. Osteocytic osteolysis: time for a second look? BoneKEy Rep. 1, 229 (2012).

37. Divieti, P. et al. Receptors for the Carboxyl-Terminal Region of PTH(1–84) Are Highly Expressed in Osteocytic Cells**This work was supported by the NIH Grant DK-11794. Endocrinology 142, 916–925 (2001).

38. Mullender, M. G., Tan, S. D., Vico, L., Alexandre, C. & Klein-Nulend, J. Differences in Osteocyte Density and Bone Histomorphometry Between Men and Women and Between Healthy and Osteoporotic Subjects. Calcif. Tissue Int. 77, 291–296 (2005).

39. Qiu, S., Rao, D. S., Palnitkar, S. & Parfitt, A. M. Reduced Iliac Cancellous Osteocyte Density in Patients With Osteoporotic Vertebral Fracture. J. Bone Miner. Res. 18, 1657– 1663 (2003).

40. Soicher, M. A., Wang, X., Zauel, R. R. & Fyhrie, D. P. Damage initiation sites in osteoporotic and normal human cancellous bone. Bone 48, 663–666 (2011).

41. Vashishth, D., Gibson, G. J. & Fyhrie, D. P. Sexual dimorphism and age dependence of osteocyte lacunar density for human vertebral cancellous bone. Anat. Rec. A. Discov. Mol. Cell. Evol. Biol. 282**A**, 157–162 (2005).

42. Qiu, S., Rao, D. S., Palnitkar, S. & Parfitt, A. M. Relationships between osteocyte density and bone formation rate in human cancellous bone. Bone 31, 709–711 (2002).

43. Aguirre, J. I. et al. Osteocyte Apoptosis Is Induced by Weightlessness in Mice and Precedes Osteoclast Recruitment and Bone Loss. J. Bone Miner. Res. 21, 605–615 (2006).

44. Webster, D. J., Morley, P. L., Van Lenthe, G. H. & Müller, R. A novel *in vivo* mouse model for mechanically stimulated bone adaptation – a combined experimental and computational validation study. Comput. Methods Biomech. Biomed. Engin. 11, 435– 441 (2008).

45. Mathavan, N. et al. Spatial transcriptomics in bone mechanomics: Exploring the mechanoregulation of fracture healing in the era of spatial omics. Sci. Adv. 11, eadp8496 (2025).

46. Bouxsein, M. L. et al. Guidelines for assessment of bone microstructure in rodents using micro–computed tomography. J. Bone Miner. Res. 25, 1468–1486 (2010).

47. Schulte, F. A., Lambers, F. M., Kuhn, G. & Müller, R. In vivo micro-computed tomography allows direct three-dimensional quantification of both bone formation and bone resorption parameters using time-lapsed imaging. Bone 48, 433–442 (2011).

48. Goff, E. et al. Large-scale quantification of human osteocyte lacunar morphological biomarkers as assessed by ultra-high-resolution desktop micro-computed tomography. Bone 152, 116094 (2021).

49. McCreadie, B. R., Hollister, S. J., Schaffler, M. B. & Goldstein, S. A. Osteocyte lacuna size and shape in women with and without osteoporotic fracture. J. Biomech. 37, 563– 572 (2004).

50. Mader, K. S., Schneider, P., Müller, R. & Stampanoni, M. A quantitative framework for the 3D characterization of the osteocyte lacunar system. Bone 57, 142–154 (2013).

51. Akhter, M. P., Kimmel, D. B., Lappe, J. M. & Recker, R. R. Effect of Macroanatomic Bone Type and Estrogen Loss on Osteocyte Lacunar Properties in Healthy Adult Women. Calcif. Tissue Int. 100, 619–630 (2017).

52. Van Der Walt, S. et al. scikit-image: image processing in Python. PeerJ 2, e453 (2014).

53. Virtanen, P. et al. SciPy 1.0: fundamental algorithms for scientific computing in Python. Nat. Methods 17, 261–272 (2020).

54. Goff, E. et al. Large-scale osteocyte lacunar morphological analysis of transiliac bone in normal and osteoporotic premenopausal women. Bone 160, 116424 (2022).

55. Seabold, S. & Perktold, J. Statsmodels: Econometric and Statistical Modeling with Python. Proc. 9th Python Sci. Conf. 2010, (2010).

