## Supplementary File for "Differential Adaptation of Osteocyte Lacunar Morphology at Bone (Re)modeling Sites under Mechanical Loading and Pharmacological Therapies in a Mouse Model of Osteoporosis"

### 1 Supplementary Material

#### 1.1 Error Quantification of the 3D-3D Multiscale Registration Workflow

The 3D-3D multiscale registration workflow involves distinct processing adaptations to register datasets from separate micro-computed tomography (micro-CT) systems. Because a true ground truth for the final high-resolution spatial (re)modeling maps was unavailable, a strict validation of the 3D-3D multiscale registration workflow was not possible. Consequently, four key image processing steps in the 3D-3D multiscale registration workflow were quantitatively evaluated through dynamic morphometric parameters (Supplementary Figure 1a). Values are presented as mean ± standard deviation (SD). Statistical difference was tested using a linear-mixed effects model with the animal as a random effect followed by Holm correction for multiple comparisons. The mismatch of (re)modeling sites on the downscaled ultra-high-resolution (ultra-HR) image was confirmed by a significant decrease in bone formation rate (BFR), bone resorption rate (BRR), mineral apposition rate (MAR), mineralizing surface (MS) and eroding surface (ES) from step 1 to step 2 (Supplementary Figure 1b-g; BFR: 1.47 ± 0.57 %/day to 0.86 ± 0.44 %/day; BRR: 0.78 ± 0.39 to 0.49 %/day ± 0.31 %/day; MAR: 1.32 ± 0.24 µm/day to 1.16 ± 0.19 µm/day; MS: 50.58 ± 9.43 % to 35.08 ± 7.77 %; ES: 27.41 ± 5.66 % to 10.96 ± 3.90 %). The mineral resorption rate (MRR) did not differ significantly between steps 1 and 2 (MRR: 1.51 ± 0.45 µm/day to 1.42 ± 0.51 µm/day). Following surface constrained projection in step 3, BFR, MS and ES increased toward their step 1 values (BFR: 0.86 ± 0.44 %/day to 1.22 ± 0.55 %/day; MS: 35.08 ± 7.78 % to 47.87 ± 6.87 %; ES: 10.97 ± 3.90 % to 19.42 ± 5.17 %), although all remained significantly different from step 1 after multiple-comparison correction. For the remaining parameters, BRR and MRR decreased further (BRR: 0.49 ± 0.31 %/day to 0.30 ± 0.12 %/day; MRR: 1.42 ± 0.51 µm/day to 0.75 ± 0.00 µm/day), whereas MAR remained statistically unchanged (1.16 ± 0.19 µm/day to 1.15 ± 0.20 µm/day). The upscaling to high resolution in step 4 introduced a significant increase in BFR (1.22 ± 0.55 %/day to 1.94 ± 0.75 %/day) suggesting an overestimation of these volumes, whereas BRR only increased slightly (0.30 ± 0.12 %/day to 0.50 ± 0.17 %/day). In contrast, MS and ES decreased significantly following upscaling (MS: 47.87 ± 6.87 % to 39.06 ± 5.59 %; ES: 19.42 ± 5.17 % to 10.18 ± 3.00 %). MAR remained statistically unchanged (1.15 ± 0.20 µm/day to 1.20 ± 0.18 µm/day), while MRR decreased significantly (0.75 ± 0.00 µm/day to 0.66 ± 0.03 µm/day). Following the surface-constrained projection, MRR exhibited a noticeable low mean value and narrow distribution. While the surface-constrained projection showed recovery of BFR, BRR, MRR as well as MS and ES, it introduced a slight reduction in the overall MAR. Given that the native nominal voxel size of the *in vivo* images is 10.5 µm, this sub-micron difference is more than an order of magnitude below the physical resolution limit of the scanner. Consequently, the difference does not compromise the size or spatial alignment of the adapted volumes. The physical resolution limit of the in vivo micro-CT furthermore explains the unique distribution pattern observed for the MRR. The MRR appears highly constrained with a remarkably narrow distribution after the surface-constrained projection. Because resorption regions are predominantly located within highly concave bone surfaces, the surface-constrained projection forces the vectors to converge. This vector crowding compresses the voxel layer, driving the MRR straight down to the absolute measurement floor of the imaging system. It is, however, not a constraining factor for the correlated lacunar analysis because resorption volumes are non-existent on the ultra-HR micro-CT images and thus essentially only the ES parameter is of interest. In addition to the surface projection, the subsequent upscaling step introduced distinct geometric effects that modulated the volume and surface metrics. While the upward-biased rounding threshold successfully restored volume-related parameters to levels comparable to *in vivo* micro-CT measurements, it resulted in a systematic decrease in the calculated contact surface area (MS and ES, Supplementary Figure 1f-g). This reduction is a predictable consequence of resolution-dependent surface mismatch. When upscaled, the formation or resorption volume cannot perfectly recapitulate the complex topographical features of the bone in the ultra-HR micro-CT. Because our downstream analysis excludes osteocyte lacunae in direct contact with the bone surface, this minor structural mismatch is negligible and does not compromise the accuracy or integrity of the regional lacunar analysis.

### 2 Supplementary Figures

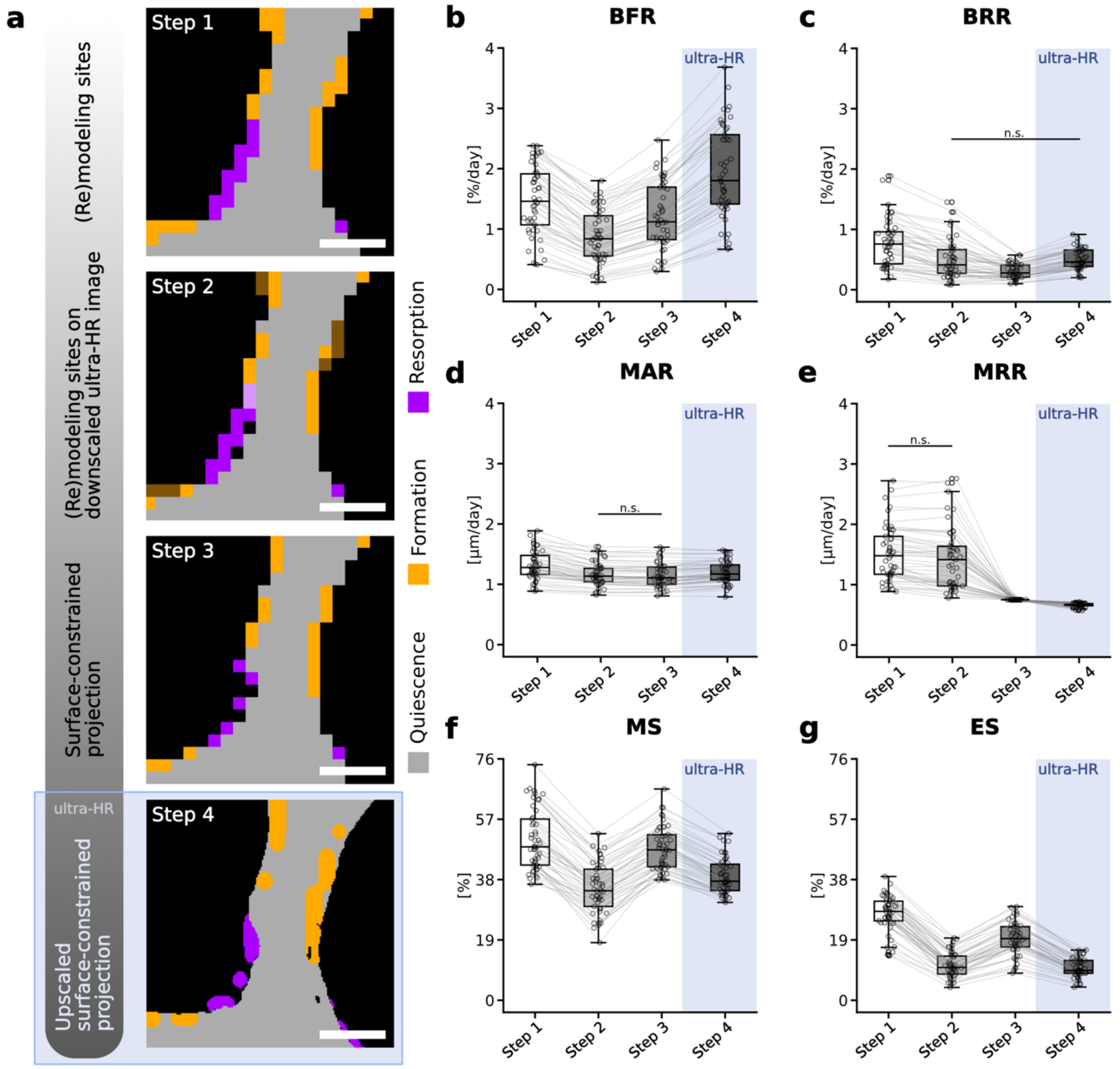


***Supplementary Figure 1: Error quantification of the 3D-3D multiscale registration workflow.*** *(a) Visual overview of the four major steps in the 3D-3D multiscale registration workflow. Step 1: in vivo (re)modeling sites, step 2: in vivo (re)modeling sites on the downscaled ultra-high resolution (ultra-HR) micro-CT, step 3: surface constrained projection of in vivo (re)modeling sites on the downscaled ultra-HR micro-CT, step 4: upscaled surface constrained projection of in vivo (re)modeling sites on original ultra-HR micro-CT. Boxplots showing (b) bone formation rate per bone volume (BFR), (c) bone resorption rate per bone volume (BRR), (d) mineral apposition rate (MAR), (e) mineral resorption rate (MRR), (f) mineralizing surface (MS) and (g) eroding surface (ES) for all steps. b-g) Each animal is shown as datapoints connected by lines to illustrate changes across workflow steps. The last step at the high-resolution of 1.2 µm voxel size is highlighted. Statistical significance was assessed using a linear-mixed effects model with animals as a random effect followed by model-based pairwise comparisons with Holm correction for multiple comparisons. Only non-significant comparisons are indicated (n.s.); comparisons not indicated were significant ($p<0.05$).*

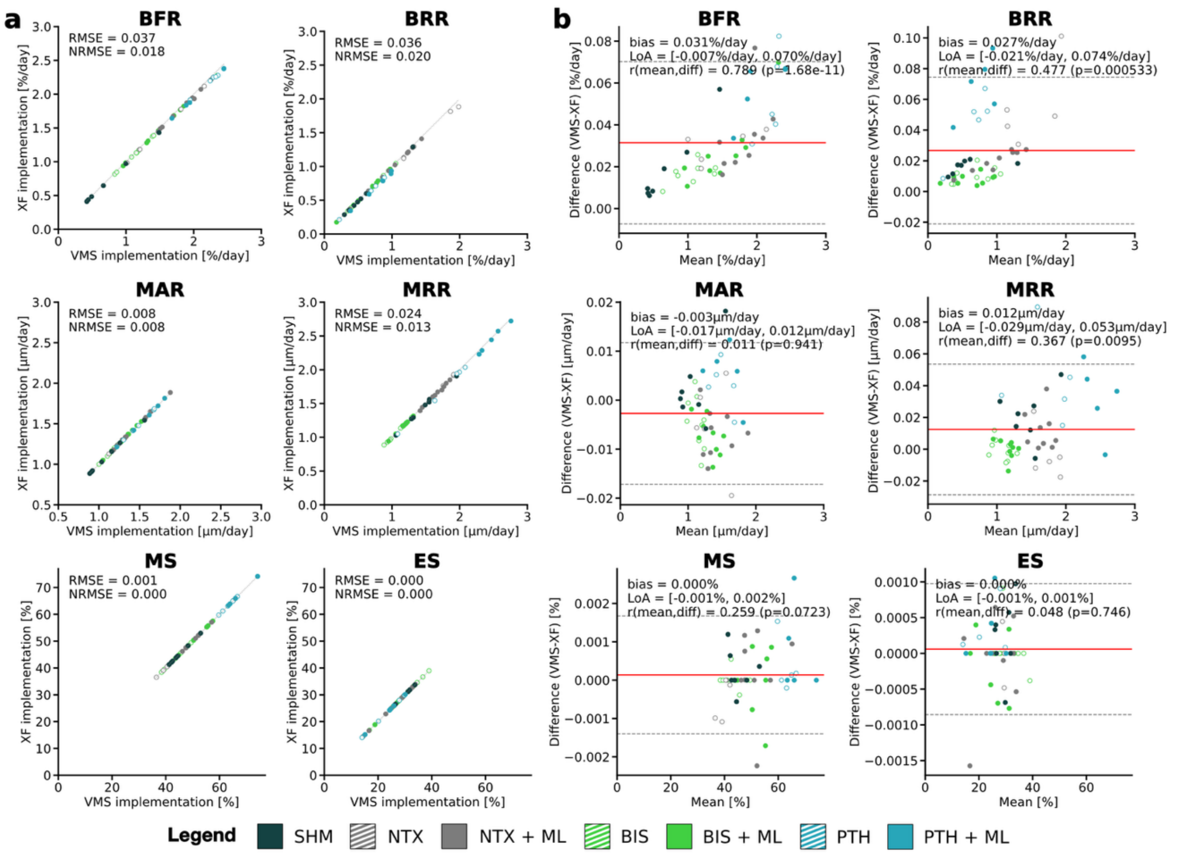


***Supplementary Figure 2:*** *(a) Dynamic morphometric parameters (bone formation rate per bone volume: BFR, bone resorption rate per bone volume: BRR, mineral apposition rate: MAR, mineral resorption rate: MRR, mineralizing surface per bone surface: MS, eroding surface per bone surface: ES) computed using the established IPL and the proposed XamFlow implementations for all samples for in vivo micro-CT scans of week 22-24. Line of equality is indicated in gray. The Root Mean Square Errors (RMSE) show good agreement with normalized RMSE (NRMSE=RMSE/($y_{max}$-$y_{min}$)) not exceeding 2%. (b) Bland-Altman plots of the dynamic morphometric parameters to assess the agreement between the two implementations. The central red line indicates mean bias and the dashed lines denote the 95% limits of agreement (LoA).*

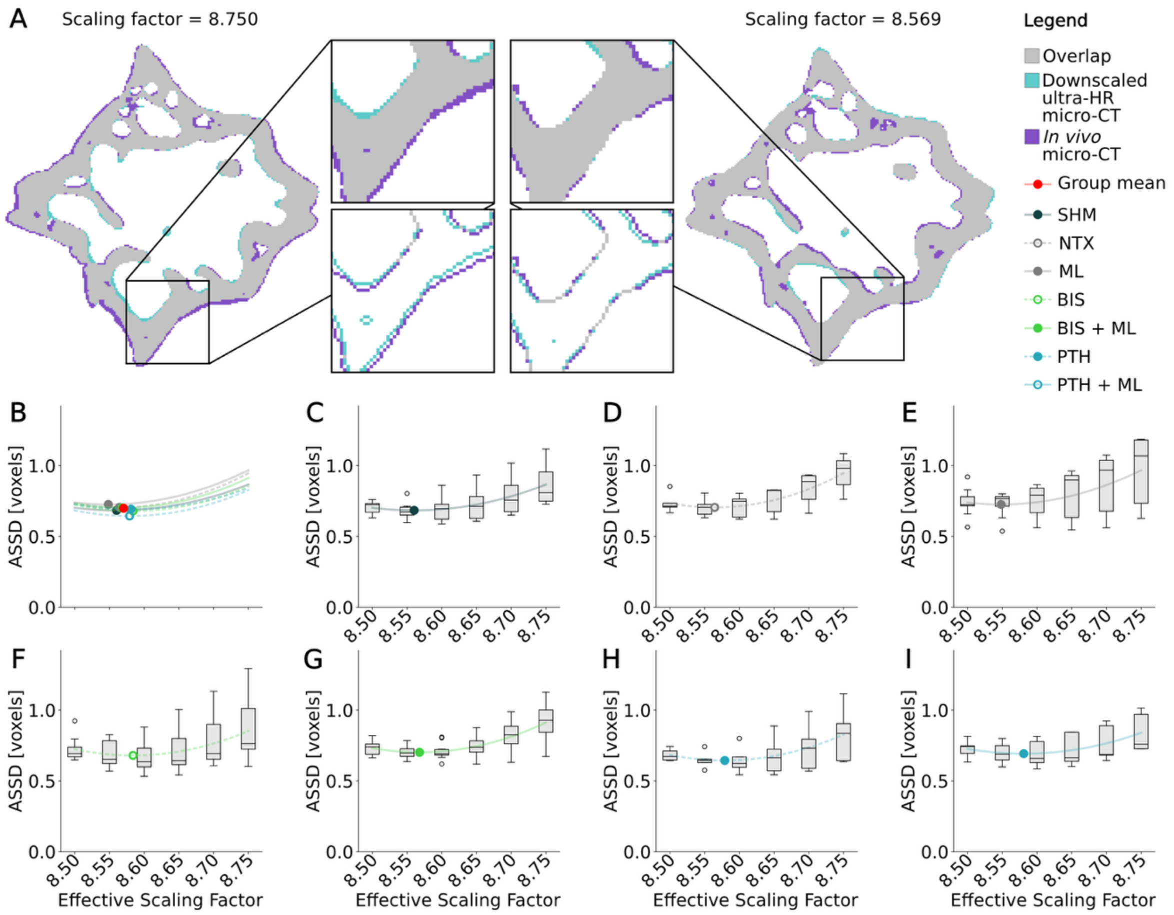


***Supplementary Figure 3:*** *Scaling factors of 8.75, 8.70, 8.65, 8.60, 8.55 and 8.50 were tested across all 50 samples. For each scaling factor, registration of the in vivo and the downscaled ultra-high-resolution (ultra-HR) micro-CT was re-run before thresholding and trabecular masking. The average symmetric surface distance (ASSD) was calculated between the trabecular bone of the thresholded resampled ultra-HR and the thresholded target in vivo micro-CT. Quadratic functions were fitted to the mean ASSD values for each scaling factor per group and the group-specific optimal scaling factor was obtained by minimizing the fitted function. The global effective scaling factor ($SF_{eff}$=8.569) was calculated as the mean of the group specific optimal scaling factors. (a) Visual representation of the relative overlap of the in vivo and the downscaled HR micro-CT at the nominal scaling factor of 8.750 and the final effective scaling factor of 8.569. (b) Fitted quadratic regression of the minimal average symmetric surface distance (ASS) between the two micro-CT images for all the groups with indicated group-specific minimum and an overall calculated mean. (c-i) ASSD for each scaling factor with fitted quadratic regression and group-specific minimum for (c) sham (SHM), (d) no treatment (NTX), (e) mechanical loading (ML), (f) bisphosphonates (BIS), (g) bisphosphonates and mechanical loading (BIS+ML), (h) parathyroid hormone (PTH) and (i) parathyroid hormone and mechanical loading (PTH+ML).*

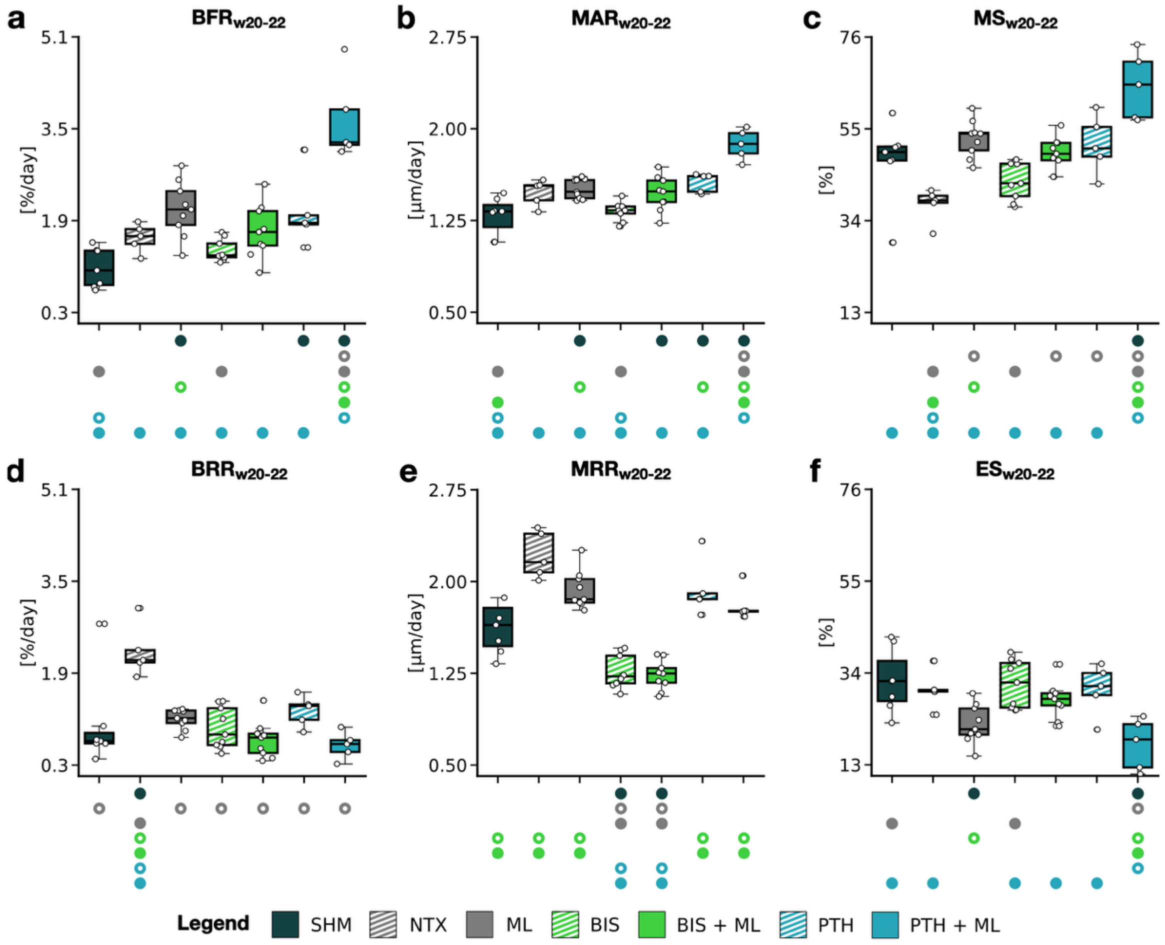


***Supplementary Figure 4:*** *Interval w20-22: Mechanical loading (ML) and pharmacological treatment, as monotreatment or combined, reverse ovariectomy induced increase in bone resorption early after the start of treatment. Boxplots of (a) bone formation rate per bone volume (BFR), (b) mineral apposition rate (MAR), (c) mineralizing surface per bone surface (MS), (d) bone resorption rate per bone volume (BRR), (e) mineral resorption rate (MRR), (f) eroded surface per bone surface (ES). Significant differences were calculated using one-way ANOVA followed by Tukey's post-hoc test. Significant differences between groups are indicated with a symbol in the color of the corresponding group (o: to monotreatment, ●: to combined treatment or sham group, see Supplementary Dataset S.1 for color encoding, p<0.05).*

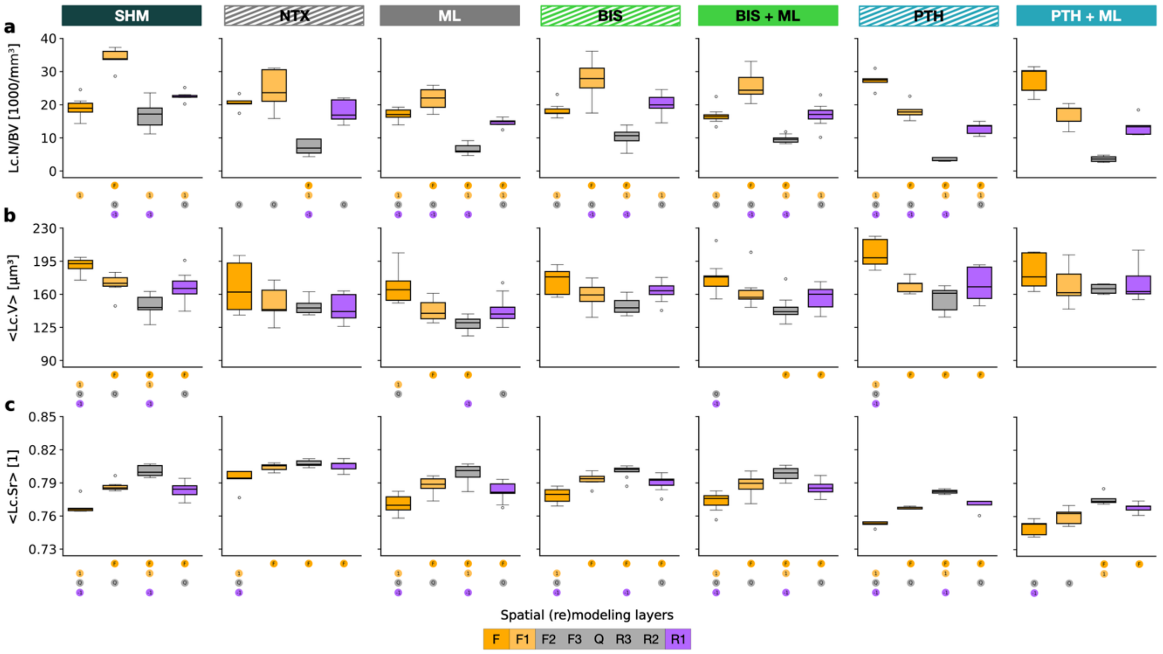


***Supplementary Figure 5:*** *Lacunar distribution and morphometry all groups between (re)modeling sites. Boxplots for (a) lacunar density (Lc.N/BV), (b) local lacunar volume (Lc.V) and (c) local lacunar sphericity (Lc.Sr) in four spatial (re)modeling regions: formation (F), formation adjacent (F1), quiescent (Q) and resorption adjacent (R1).*

### 3 Supplementary Tables

***Supplementary Table 1:*** *Quadratic functions fitted to the group specific mean average symmetric surface distance (ASSD) per scaling factor. The minimum of each function ($y_{min}$ and $x_{min}$) was extracted, where $y_{min}$ is the minimum ASSD [voxels] and $x_{min}$ is the corresponding group-specific scaling factor.*

| **Group** | **Fitted quadratic function** | $\mathbf{y_{min}}$ | $\mathbf{x_{min}}$ |
|---|---|---|---|
| SHM | $y = 5.071x^2 - 86.817x + 372.270$ | 0.684 | 8.560 |
| NTX | $y = 7.003x^2 - 119.946x + 514.349$ | 0.705 | 8.564 |
| ML | $y = 5.932x^2 - 101.420x + 434.212$ | 0.726 | 8.548 |
| BIS | $y = 6.262x^2 - 107.508x + 462.099$ | 0.680 | 8.584 |
| BIS+ML | $y = 6.332x^2 - 108.498x + 465.501$ | 0.702 | 8.568 |
| PTH | $y = 6.232x^2 - 106.923x + 459.286$ | 0.644 | 8.579 |
| PTH+ML | $y = 5.160x^2 - 88.559x + 380.654$ | 0.692 | 8.581 |
| | **Mean:** | **0.690** | **8.569** |

***Supplementary Table 2:*** *Color encoding for significant differences to a specific group. Significances are depicted with a symbol in the color of the corresponding group using two-way ANOVA followed by Tukey's post-hoc test (p<0.05:* ***o:*** *to monotreatment, ●: to combined treatment or sham group, ●●o●o●o: to SHM, to NTX, to ML, to BIS, to BIS+ML, to PTH, to PTH+ML).*

| ● | ● | o | ● | o | ● | o |
|---|---|---|---|---|---|---|
| SHM | NTX | ML | BIS | BIS+ML | PTH | PTH+ML |

***Supplementary Table 3:*** *Static morphometric parameters of the trabecular compartment of non-loaded (0N) and loaded (8N) 6th caudal vertebrae at weeks 15, 20, 22 and 24 (w15, w20, w22, w24). Values are presented as group means ± standard error, Δ represents the %-change from w20 to w24. One-way ANOVA for the shown groups followed by Tukey's post-hoc test was used to test differences between groups. 'X' denotes the reference group. Significant differences are indicated with (*) p<0.05, (**) p<0.01, (***) p<0.001.*

| Readout | ML | w15 | **w20 (baseline)** | w22 | w24 | **Δw20-24 [%]** | SHM ● | NTX ○ | ML ● | BIS ○ | BIS+ML ● | PTH ○ | PTH+ML ● |
|---|---|---|---|---|---|---|---|---|---|---|---|---|---|
| **SHM (0N: n=7)** | | | | | | | | | | | | | |
| BV/TV [%] | 0N | 17.382 ± 0.871 | 20.266 ± 1.015 | 20.709 ± 1.439 | 21.481 ± 1.479 | 6.026 ± 5.070 | X | 0.789 | ** 0.001 | 0.719 | ** 0.004 | *** 0.001 | *** 0.000 |
| BS/TV [%] | 0N | 6.909 ± 0.269 | 6.667 ± 0.253 | 6.558 ± 0.297 | 6.507 ± 0.299 | -2.392 ± 2.494 | X | 0.118 | 0.979 | 0.053 | 0.365 | ** 0.004 | *** 0.000 |
| Tb.Th [µm] | 0N | 0.067 ± 0.001 | 0.078 ± 0.002 | 0.080 ± 0.002 | 0.083 ± 0.002 | 6.065 ± 2.049 | X | 1.000 | *** 0.000 | 0.999 | *** 0.000 | 0.119 | *** 0.000 |
| Tb.N [1/mm] | 0N | 3.172 ± 0.097 | 3.025 ± 0.103 | 2.966 ± 0.097 | 2.943 ± 0.093 | -2.568 ± 1.426 | X | * 0.038 | ** 0.002 | 0.390 | 0.993 | 1.000 | 0.997 |
| **NTX (0N: n=5) and ML (8N: n=9)** | | | | | | | | | | | | | |
| BV/TV [%] | 0N | 18.735 ± 2.008 | 12.578 ± 0.252 | 11.740 ± 0.172 | 12.178 ± 0.186 | -2.972 ± 2.851 | 0.789 | X | *** 0.000 | 0.083 | *** 0.000 | *** 0.000 | *** 0.000 |
| | 8N | 19.084 ± 0.640 | 13.945 ± 0.648 | 16.025 ± 0.603 | 17.999 ± 0.723 | 30.069 ± 4.997 | ** 0.001 | *** 0.000 | X | 0.055 | 1.000 | 0.966 | *** 0.000 |
| BS/TV [%] | 0N | 6.966 ± 0.520 | 5.828 ± 0.245 | 5.315 ± 0.221 | 5.284 ± 0.207 | -9.212 ± 5.385 | 0.118 | X | 0.374 | *** 0.000 | *** 0.000 | *** 0.000 | *** 0.000 |
| | 8N | 7.040 ± 0.189 | 6.133 ± 0.213 | 5.998 ± 0.202 | 5.869 ± 0.212 | -4.209 ± 5.884 | 0.979 | 0.374 | X | ** 0.003 | * 0.043 | *** 0.000 | *** 0.000 |
| Tb.Th [µm] | 0N | 0.070 ± 0.002 | 0.061 ± 0.003 | 0.062 ± 0.002 | 0.065 ± 0.002 | 5.215 ± 1.530 | 1.000 | X | *** 0.000 | 1.000 | *** 0.001 | 0.108 | *** 0.000 |
| | 8N | 0.071 ± 0.002 | 0.064 ± 0.002 | 0.072 ± 0.001 | 0.081 ± 0.002 | 28.003 ± 2.678 | *** 0.000 | *** 0.000 | X | *** 0.000 | * 0.025 | ** 0.002 | 0.471 |
| Tb.N [1/mm] | 0N | 3.213 ± 0.153 | 2.860 ± 0.103 | 2.669 ± 0.081 | 2.596 ± 0.087 | -9.162 ± 1.251 | * 0.038 | X | 0.999 | *** 0.000 | ** 0.005 | * 0.029 | * 0.018 |
| | 8N | 3.245 ± 0.084 | 3.012 ± 0.101 | 2.838 ± 0.100 | 2.708 ± 0.091 | -10.064 ± 1.163 | ** 0.002 | 0.999 | X | *** 0.000 | *** 0.000 | ** 0.002 | ** 0.001 |

**BIS (0N: n=9, 8N: n=9)**

| | | | | | | | | | | | | | |
|---|---|---|---|---|---|---|---|---|---|---|---|---|---|
| BV/TV [%] | 0N | 17.249 ± 0.844 | 13.119 ± 0.381 | 13.965 ± 0.209 | 14.942 ± 0.224 | 14.448 ± 2.834 | 0.719 | 0.997 | 0.000 | X | 0.134 | * 0.017 | *** 0.000 |
| | 8N | 18.046 ± 0.800 | 14.315 ± 0.538 | 15.827 ± 0.506 | 18.287 ± 0.564 | 28.040 ± 1.483 | ** 0.004 | *** 0.000 | 1.000 | 0.134 | X | 0.865 | *** 0.000 |
| BS/TV [%] | 0N | 6.591 ± 0.241 | 5.858 ± 0.144 | 5.929 ± 0.131 | 6.101 ± 0.131 | 4.422 ± 6.151 | 0.053 | *** 0.000 | ** 0.003 | X | 0.945 | 0.772 | 0.258 |
| | 8N | 6.498 ± 0.236 | 6.073 ± 0.211 | 6.058 ± 0.208 | 6.195 ± 0.184 | 2.162 ± 6.092 | 0.365 | *** 0.000 | * 0.043 | 0.945 | X | * 0.040 | 0.365 |
| Tb.Th [µm] | 0N | 0.069 ± 0.002 | 0.063 ± 0.001 | 0.065 ± 0.001 | 0.067 ± 0.001 | 4.832 ± 1.334 | 0.999 | 1.000 | *** 0.000 | X | *** 0.000 | * 0.034 | *** 0.000 |
| | 8N | 0.073 ± 0.002 | 0.066 ± 0.001 | 0.071 ± 0.001 | 0.078 ± 0.002 | 19.271 ± 0.832 | *** 0.001 | *** 0.001 | * 0.025 | *** 0.000 | X | 0.764 | *** 0.000 |
| Tb.N [1/mm] | 0N | 3.049 ± 0.091 | 2.900 ± 0.087 | 2.890 ± 0.082 | 2.929 ± 0.079 | 0.429 ± 0.801 | 0.390 | *** 0.000 | *** 0.000 | X | 0.753 | 0.758 | 0.861 |
| | 8N | 3.002 ± 0.075 | 2.928 ± 0.077 | 2.883 ± 0.086 | 2.887 ± 0.070 | -1.365 ± 0.722 | 0.993 | ** 0.005 | *** 0.000 | 0.753 | X | 1.000 | 1.000 |

**PTH (0N: n=5, 8N: n=5)**

| | | | | | | | | | | | | | |
|---|---|---|---|---|---|---|---|---|---|---|---|---|---|
| BV/TV [%] | 0N | 18.838 ± 1.465 | 14.753 ± 0.818 | 16.921 ± 0.737 | 19.941 ± 0.987 | 35.678 ± 4.619 | *** 0.001 | *** 0.000 | 0.966 | * 0.017 | 0.865 | X | *** 0.001 |
| | 8N | 18.730 ± 1.195 | 14.007 ± 0.797 | 19.037 ± 0.716 | 23.244 ± 1.132 | 66.741 ± 6.052 | *** 0.000 | *** 0.000 | *** 0.000 | *** 0.000 | *** 0.000 | *** 0.001 | X |
| BS/TV [%] | 0N | 6.348 ± 0.235 | 5.995 ± 0.179 | 6.320 ± 0.165 | 6.455 ± 0.199 | 7.691 ± 6.645 | ** 0.004 | *** 0.000 | *** 0.000 | 0.772 | * 0.040 | X | 0.987 |
| | 8N | 6.645 ± 0.232 | 6.239 ± 0.270 | 6.439 ± 0.249 | 6.842 ± 0.270 | 9.772 ± 7.151 | *** 0.000 | *** 0.000 | *** 0.000 | 0.258 | 0.365 | 0.987 | X |
| Tb.Th [µm] | 0N | 0.077 ± 0.004 | 0.067 ± 0.002 | 0.069 ± 0.002 | 0.077 ± 0.002 | 14.794 ± 1.926 | 0.119 | 0.108 | ** 0.002 | * 0.034 | *** 0.000 | X | *** 0.000 |
| | 8N | 0.073 ± 0.002 | 0.063 ± 0.001 | 0.075 ± 0.002 | 0.084 ± 0.002 | 33.917 ± 3.692 | *** 0.000 | *** 0.000 | 0.471 | 0.764 | *** 0.000 | *** 0.000 | X |
| Tb.N [1/mm] | 0N | 2.995 ± 0.090 | 2.880 ± 0.077 | 2.828 ± 0.085 | 2.829 ± 0.080 | -1.796 ± 0.597 | 1.000 | * 0.029 | ** 0.002 | 0.758 | 1.000 | X | 1.000 |
| | 8N | 3.137 ± 0.122 | 3.109 ± 0.128 | 3.003 ± 0.138 | 3.068 ± 0.172 | -1.401 ± 3.097 | 0.997 | * 0.018 | ** 0.001 | 0.861 | 1.000 | 1.000 | X |

***Supplementary Table 4:*** *Dynamic morphometric parameters of the trabecular compartment of non-loaded (0N) and loaded (8N) 6th caudal vertebrae at weeks 15, 20, 22 and 24 (w15, w20, w22, w24). Values are presented as group means ± standard error in week intervals w20-22 and w22-24. The %-change between w20-22 and w22-24, calculated per animal, is depicted as well. Two-way ANOVA over time for the shown groups followed by Tukey's post-hoc test was used to test differences between groups. 'X' denotes the reference group. Significant differences are indicated with (*) p<0.05, (**) p<0.01, (***) p<0.001.*

| Readout | N | **w20-22** | **w22-24** | **%-change between w20-22 and w22-24 [%]** | | SHM ● | NTX ○ | ML ● | BIS ○ | BIS+ML ● | PTH ○ | PTH+ML ● |
|---|---|---|---|---|---|---|---|---|---|---|---|---|
| **SHM (0N: n=7)** | | | | | | | | | | | | |
| BFR [%/d] | 0N | 1.079 ± 0.130 | 0.704 ± 0.156 | -26.071 ± 20.228 | w20-22 | X | 0.460 | ** 0.001 | 0.853 | 0.106 | * 0.012 | *** 0.000 |
| | | | | | w22-24 | X | * 0.013 | *** 0.000 | 0.602 | *** 0.000 | *** 0.000 | *** 0.000 |
| BRR [%/d] | 0N | 0.995 ± 0.301 | 0.549 ± 0.101 | -35.079 ± 9.249 | w20-22 | X | 0.113 | 0.935 | ** 0.005 | ** 0.003 | 0.965 | 1.000 |
| | | | | | w22-24 | X | *** 0.000 | 0.479 | 1.000 | 1.000 | 1.000 | 1.000 |
| MAR [µm/d] | 0N | 1.288 ± 0.060 | 1.097 ± 0.081 | -14.643 ± 4.975 | w20-22 | X | 0.087 | ** 0.007 | 0.995 | * 0.043 | ** 0.003 | *** 0.000 |
| | | | | | w22-24 | X | 0.157 | * 0.017 | 0.991 | 0.056 | *** 0.000 | *** 0.001 |
| MRR [µm/d] | 0N | 1.791 ± 0.220 | 1.340 ± 0.104 | -23.320 ± 3.097 | w20-22 | X | 0.113 | 0.935 | ** 0.005 | ** 0.003 | 0.965 | 1.000 |
| | | | | | w22-24 | X | 0.091 | * 0.015 | 0.169 | 0.708 | *** 0.001 | *** 0.000 |
| MS [%] | 0N | 47.811 ± 3.428 | 43.942 ± 1.177 | -3.446 ± 11.592 | w20-22 | X | 0.059 | 0.619 | 0.687 | 0.999 | 0.935 | *** 0.000 |
| | | | | | w22-24 | X | 0.876 | *** 0.000 | 0.984 | *** 0.000 | *** 0.000 | *** 0.000 |
| ES [%] | 0N | 32.241 ± 2.761 | 30.141 ± 1.062 | -2.110 ± 9.149 | w20-22 | X | 0.994 | * 0.011 | 1.000 | 0.656 | 0.995 | *** 0.001 |
| | | | | | w22-24 | X | 1.000 | * 0.027 | 1.000 | 0.458 | * 0.022 | ** 0.458 |

| | | | | | | | | | | | | |
|---|---|---|---|---|---|---|---|---|---|---|---|---|
| **NTX (0N: n=5) and ML (8N: n=9)** | | | | | | | | | | | | |
| BFR [%/d] | 0N | 1.600 ± 0.110 | 1.385 ± 0.192 | -11.451 ± 10.866 | w20-22 | 0.460 | X | 0.454 | 0.973 | 0.999 | 0.705 | *** 0.000 |
| | | | | | w22-24 | * 0.013 | X | 0.840 | 0.301 | 0.997 | * 0.023 | 0.053 |
| | 8N | 2.099 ± 0.163 | 1.622 ± 0.083 | -5.716 ± 4.788 | w20-22 | ** 0.001 | 0.454 | X | * 0.025 | 0.568 | 1.000 | *** 0.000 |
| | | | | | w22-24 | *** 0.000 | 0.840 | X | ** 0.002 | 0.978 | 0.173 | 0.334 |
| BRR [%/d] | 0N | 2.277 ± 0.203 | 1.297 ± 0.145 | -43.464 ± 2.382 | w20-22 | 0.113 | X | 0.482 | *** 0.000 | *** 0.000 | 0.635 | 0.209 |
| | | | | | w22-24 | *** 0.000 | X | ** 0.006 | *** 0.000 | *** 0.000 | *** 0.000 | *** 0.001 |
| | 8N | 1.102 ± 0.058 | 0.778 ± 0.081 | -29.893 ± 5.940 | w20-22 | 0.935 | 0.482 | X | *** 0.000 | *** 0.000 | 1.000 | 0.978 |
| | | | | | w22-24 | 0.479 | ** 0.006 | X | 0.641 | 0.563 | 0.756 | 0.854 |
| MAR [µm/d] | 0N | 1.478 ± 0.048 | 1.324 ± 0.077 | -10.067 ± 5.648 | w20-22 | 0.087 | X | 0.999 | 0.214 | 1.000 | 0.908 | *** 0.000 |
| | | | | | w22-24 | 0.157 | X | 0.999 | 0.395 | 1.000 | 0.346 | 0.451 |
| | 8N | 1.508 ± 0.027 | 1.364 ± 0.048 | -9.491 ± 2.893 | w20-22 | ** 0.007 | 0.999 | X | * 0.021 | 0.990 | 0.980 | *** 0.000 |
| | | | | | w22-24 | * 0.017 | 0.999 | X | 0.063 | 0.999 | 0.455 | 0.584 |
| MRR [µm/d] | 0N | 2.215 ± 0.086 | 1.647 ± 0.076 | -25.710 ± 1.400 | w20-22 | 0.113 | X | 0.482 | *** 0.000 | *** 0.000 | 0.635 | 0.209 |
| | | | | | w22-24 | 0.091 | X | 1.000 | *** 0.000 | ** 0.001 | 0.594 | *** 0.000 |
| | 8N | 1.933 ± 0.052 | 1.670 ± 0.050 | -13.462 ± 2.147 | w20-22 | 0.935 | 0.482 | X | *** 0.000 | *** 0.000 | 0.694 | 0.973 |
| | | | | | w22-24 | * 0.015 | 1.000 | X | *** 0.000 | *** 0.000 | * 0.026 | 0.301 |
| MS [%] | 0N | 37.665 ± 1.734 | 41.157 ± 0.502 | 10.080 ± 4.436 | w20-22 | 0.059 | X | *** 0.001 | 0.597 | * 0.012 | ** 0.008 | *** 0.000 |
| | | | | | w22-24 | 0.876 | X | *** 0.000 | 0.997 | *** 0.000 | *** 0.000 | *** 0.000 |

| | | | | | | | | | | | | |
|---|---|---|---|---|---|---|---|---|---|---|---|---|
| | 8N | 52.723 ± 1.415 | 56.544 ± 1.876 | 7.318 ± 2.417 | w20-22 | 0.619 | *** 0.001 | X | * 0.017 | 0.859 | ** 0.008 | *** 0.000 |
| | | | | | w22-24 | *** 0.000 | *** 0.000 | X | *** 0.000 | 0.535 | *** 0.000 | *** 0.000 |
| ES [%] | 0N | 30.180 ± 1.955 | 30.194 ± 0.444 | 1.459 ± 5.604 | w20-22 | 0.994 | X | 0.140 | 1.000 | 0.984 | 1.000 | * 0.011 |
| | | | | | w22-24 | 1.000 | X | 0.057 | 1.000 | 0.563 | * 0.040 | *** 0.000 |
| | 8N | 22.285 ± 1.490 | 23.272 ± 1.757 | 5.630 ± 7.072 | w20-22 | * 0.011 | 0.140 | X | * 0.013 | 0.328 | 0.136 | 0.724 |
| | | | | | w22-24 | * 0.027 | 0.057 | X | * 0.037 | 0.754 | 0.997 | 0.964 |
| **BIS (0N: n=9, 8N: n=9)** | | | | | | | | | | | | |
| BFR [%/d] | 0N | 1.375 ± 0.063 | 0.984 ± 0.083 | -24.165 ± 6.494 | w20-22 | 0.853 | 0.973 | * 0.025 | X | 0.694 | 0.135 | *** 0.000 |
| | | | | | w22-24 | 0.602 | 0.301 | ** 0.002 | X | * 0.026 | *** 0.000 | *** 0.000 |
| | 8N | 1.716 ± 0.156 | 1.493 ± 0.124 | -0.886 ± 6.565 | w20-22 | 0.106 | 0.999 | 0.568 | 0.694 | X | 0.839 | *** 0.000 |
| | | | | | w22-24 | *** 0.000 | 0.997 | 0.978 | * 0.026 | X | * 0.034 | 0.082 |
| BRR [%/d] | 0N | 0.947 ± 0.120 | 0.592 ± 0.079 | -36.308 ± 5.376 | w20-22 | ** 0.005 | *** 0.000 | *** 0.000 | X | 1.000 | *** 0.001 | * 0.010 |
| | | | | | w22-24 | 1.000 | *** 0.000 | 0.641 | X | 1.000 | 1.000 | 1.000 |
| | 8N | 0.745 ± 0.108 | 0.578 ± 0.068 | -18.239 ± 7.643 | w20-22 | ** 0.003 | *** 0.000 | *** 0.000 | 1.000 | X | *** 0.000 | ** 0.006 |
| | | | | | w22-24 | 1.000 | *** 0.000 | 0.563 | 1.000 | X | 1.000 | 1.000 |
| MAR [µm/d] | 0N | 1.325 ± 0.025 | 1.151 ± 0.033 | -12.895 ± 2.740 | w20-22 | 0.995 | 0.214 | * 0.021 | X | 0.124 | ** 0.009 | *** 0.000 |
| | | | | | w22-24 | 0.991 | 0.395 | 0.063 | X | 0.191 | ** 0.001 | ** 0.002 |
| | 8N | 1.469 ± 0.048 | 1.328 ± 0.041 | -9.312 ± 2.371 | w20-22 | * 0.043 | 1.000 | 0.990 | 0.124 | X | 0.771 | *** 0.000 |
| | | | | | w22-24 | 0.056 | 1.000 | 0.999 | 0.191 | X | 0.227 | 0.323 |

| | | | | | | | | | | | | |
|---|---|---|---|---|---|---|---|---|---|---|---|---|
| | 8N | 1.234 ± 0.041 | 1.195 ± 0.019 | -2.511 ± 2.853 | w20-22 | ** 0.003 | *** 0.000 | *** 0.000 | 1.000 | X | *** 0.000 | ** 0.006 |
| | | | | | w22-24 | 0.708 | ** 0.001 | *** 0.000 | 0.937 | X | *** 0.000 | *** 0.000 |
| MS [%] | 0N | 43.210 ± 1.460 | 42.389 ± 1.092 | -1.195 ± 3.476 | w20-22 | 0.687 | 0.597 | * 0.017 | X | 0.293 | 0.165 | *** 0.000 |
| | | | | | w22-24 | 0.984 | 0.997 | *** 0.000 | X | *** 0.000 | *** 0.000 | *** 0.000 |
| | 8N | 49.278 ± 1.266 | 53.222 ± 1.160 | 8.439 ± 3.108 | w20-22 | 0.999 | * 0.012 | 0.859 | 0.293 | X | 0.994 | *** 0.000 |
| | | | | | w22-24 | *** 0.000 | *** 0.000 | 0.535 | *** 0.000 | X | *** 0.000 | *** 0.000 |
| ES [%] | 0N | 31.464 ± 1.830 | 29.459 ± 1.287 | -5.278 ± 3.625 | w20-22 | 1.000 | 1.000 | * 0.013 | X | 0.772 | 1.000 | *** 0.001 |
| | | | | | w22-24 | 1.000 | 1.000 | * 0.037 | X | 0.598 | * 0.030 | * 0.012 |
| | 8N | 27.798 ± 1.385 | 26.123 ± 1.562 | -5.820 ± 4.220 | w20-22 | 0.656 | 0.984 | 0.328 | 0.772 | X | 0.982 | * 0.025 |
| | | | | | w22-24 | 0.458 | 0.563 | 0.754 | 0.598 | X | 0.537 | 0.322 |
| **PTH (0N: n=5, 8N: n=5)** | | | | | | | | | | | | |
| BFR [%/d] | 0N | 2.051 ± 0.287 | 2.077 ± 0.110 | 17.820 ± 14.319 | w20-22 | * 0.012 | 0.705 | 1.000 | 0.135 | 0.839 | X | *** 0.000 |
| | | | | | w22-24 | *** 0.000 | * 0.023 | 0.173 | *** 0.000 | * 0.034 | X | 1.000 |
| | 8N | 3.661 ± 0.331 | 2.011 ± 0.113 | -25.050 ± 5.085 | w20-22 | *** 0.000 | *** 0.000 | *** 0.000 | *** 0.000 | *** 0.000 | *** 0.000 | X |
| | | | | | w22-24 | *** 0.000 | 0.053 | 0.334 | *** 0.000 | 0.082 | 1.000 | X |
| BRR [%/d] | 0N | 1.244 ± 0.119 | 0.583 ± 0.092 | -52.696 ± 7.464 | w20-22 | 0.965 | 0.635 | 1.000 | *** 0.001 | *** 0.000 | X | 0.987 |
| | | | | | w22-24 | 1.000 | *** 0.000 | 0.756 | 1.000 | 1.000 | X | 1.000 |
| | 8N | 0.643 ± 0.107 | 0.608 ± 0.066 | 2.098 ± 14.191 | w20-22 | 1.000 | 0.209 | 0.978 | * 0.010 | ** 0.006 | 0.987 | X |
| | | | | | w22-24 | 1.000 | *** 0.001 | 0.854 | 1.000 | 1.000 | 1.000 | X |

| | | | | | | | | | | | | |
|---|---|---|---|---|---|---|---|---|---|---|---|---|
| MAR [µm/d] | 0N | 1.560 ± 0.035 | 1.528 ± 0.076 | -2.095 ± 4.309 | w20-22 | ** 0.003 | 0.908 | 0.980 | ** 0.009 | 0.771 | X | ** 0.002 |
| | | | | | w22-24 | *** 0.000 | 0.346 | 0.455 | ** 0.001 | 0.227 | X | 1.000 |
| | 8N | 1.872 ± 0.056 | 1.511 ± 0.058 | -19.346 ± 1.034 | w20-22 | *** 0.000 | *** 0.000 | *** 0.000 | *** 0.000 | *** 0.000 | ** 0.002 | X |
| | | | | | w22-24 | *** 0.001 | 0.451 | 0.584 | ** 0.002 | 0.323 | 1.000 | X |
| MRR [µm/d] | 0N | 1.935 ± 0.103 | 1.849 ± 0.117 | -4.229 ± 5.146 | w20-22 | 0.965 | 0.635 | 0.694 | *** 0.001 | *** 0.000 | X | 0.987 |
| | | | | | w22-24 | *** 0.001 | 0.594 | * 0.026 | *** 0.000 | *** 0.000 | X | *** 0.000 |
| | 8N | 1.808 ± 0.061 | 2.460 ± 0.106 | 36.409 ± 6.309 | w20-22 | 1.000 | 0.209 | 0.973 | * 0.010 | ** 0.006 | 0.987 | X |
| | | | | | w22-24 | *** 0.000 | *** 0.000 | 0.301 | *** 0.000 | *** 0.000 | *** 0.000 | X |
| MS [%] | 0N | 51.385 ± 2.983 | 64.397 ± 1.665 | 26.392 ± 4.876 | w20-22 | 0.935 | ** 0.008 | ** 0.008 | 0.165 | 0.994 | X | ** 0.010 |
| | | | | | w22-24 | *** 0.000 | *** 0.000 | *** 0.000 | *** 0.000 | *** 0.000 | X | 0.778 |
| | 8N | 64.880 ± 3.414 | 67.895 ± 1.820 | 5.507 ± 4.838 | w20-22 | *** 0.000 | *** 0.000 | *** 0.000 | *** 0.000 | *** 0.000 | ** 0.010 | X |
| | | | | | w22-24 | *** 0.000 | *** 0.000 | *** 0.000 | *** 0.000 | *** 0.000 | 0.778 | X |
| ES [%] | 0N | 30.228 ± 2.598 | 21.962 ± 2.312 | -26.350 ± 7.426 | w20-22 | 0.995 | 1.000 | 0.136 | 1.000 | 0.982 | X | * 0.011 |
| | | | | | w22-24 | * 0.022 | * 0.040 | 0.997 | * 0.030 | 0.537 | X | 1.000 |
| | 8N | 17.698 ± 2.628 | 21.138 ± 1.091 | 29.969 ± 18.499 | w20-22 | *** 0.001 | * 0.011 | 0.724 | *** 0.001 | * 0.025 | * 0.011 | X |
| | | | | | w22-24 | ** 0.458 | *** 0.000 | 0.964 | * 0.012 | 0.322 | 1.000 | X |
